# Atrial Proteomic Profiling Reveals a Switch Towards Profibrotic Gene Expression Program in CREM-IbΔC-X Mice with Persistent Atrial Fibrillation

**DOI:** 10.1101/2024.01.10.575097

**Authors:** Shuai Zhao, Mohit M. Hulsurkar, Satadru K. Lahiri, Yuriana Aguilar-Sanchez, Elda Munivez, Frank Ulrich Müller, Antrix Jain, Anna Malovannaya, Kendrick Yiu, Svetlana Reilly, Xander H.T. Wehrens

## Abstract

**Background:** Overexpression of the CREM (cAMP response element-binding modulator) isoform CREM-IbΔC-X in transgenic mice (CREM-Tg) causes the age-dependent development of spontaneous AF.

**Purpose:** To identify key proteome signatures and biological processes accompanying the development of persistent AF through integrated proteomics and bioinformatics analysis.

**Methods:** Atrial tissue samples from three CREM-Tg mice and three wild-type littermates were subjected to unbiased mass spectrometry-based quantitative proteomics, differential expression and pathway enrichment analysis, and protein-protein interaction (PPI) network analysis.

**Results:** A total of 98 differentially expressed proteins were identified. Gene ontology analysis revealed enrichment for biological processes regulating actin cytoskeleton organization and extracellular matrix (ECM) dynamics. Changes in ITGAV, FBLN5, and LCP1 were identified as being relevant to atrial fibrosis and remodeling based on expression changes, co-expression patterns, and PPI network analysis. Comparative analysis with previously published datasets revealed a shift in protein expression patterns from ion-channel and metabolic regulators in young CREM-Tg mice to profibrotic remodeling factors in older CREM-Tg mice. Furthermore, older CREM-Tg mice exhibited protein expression patterns that resembled those of humans with persistent AF.

**Conclusions:** This study uncovered distinct temporal changes in atrial protein expression patterns with age in CREM-Tg mice consistent with the progressive evolution of AF. Future studies into the role of the key differentially abundant proteins identified in this study in AF progression may open new therapeutic avenues to control atrial fibrosis and substrate development in AF.

**Graphical abstract:** Graphical abstract summarizing key findings of this paper. The atrial proteome in 9-month-old CREM- Tg mice with chronic persistent AF (perAF) was compared with age-matched WT littermates. In addition, proteome changes in these old CREM-Tg mice were compared with proteome changes previously identified in young CREM-Tg mice with paroxysmal AF (pAF). Moreover, an interspecies comparison was performed between old CREM-Tg mice and human patients with perAF. The major findings are that in pAF, key changes were identified in proteins involved in metabolism, energy production, DNA synthesis, and cell proliferation and growth. On the other hand, in mice and humans with perAF, key changes were found in the expression of proteins involved in collagen production, extracellular matrix remodeling, actin cytoskeleton organization, and tissue repair.

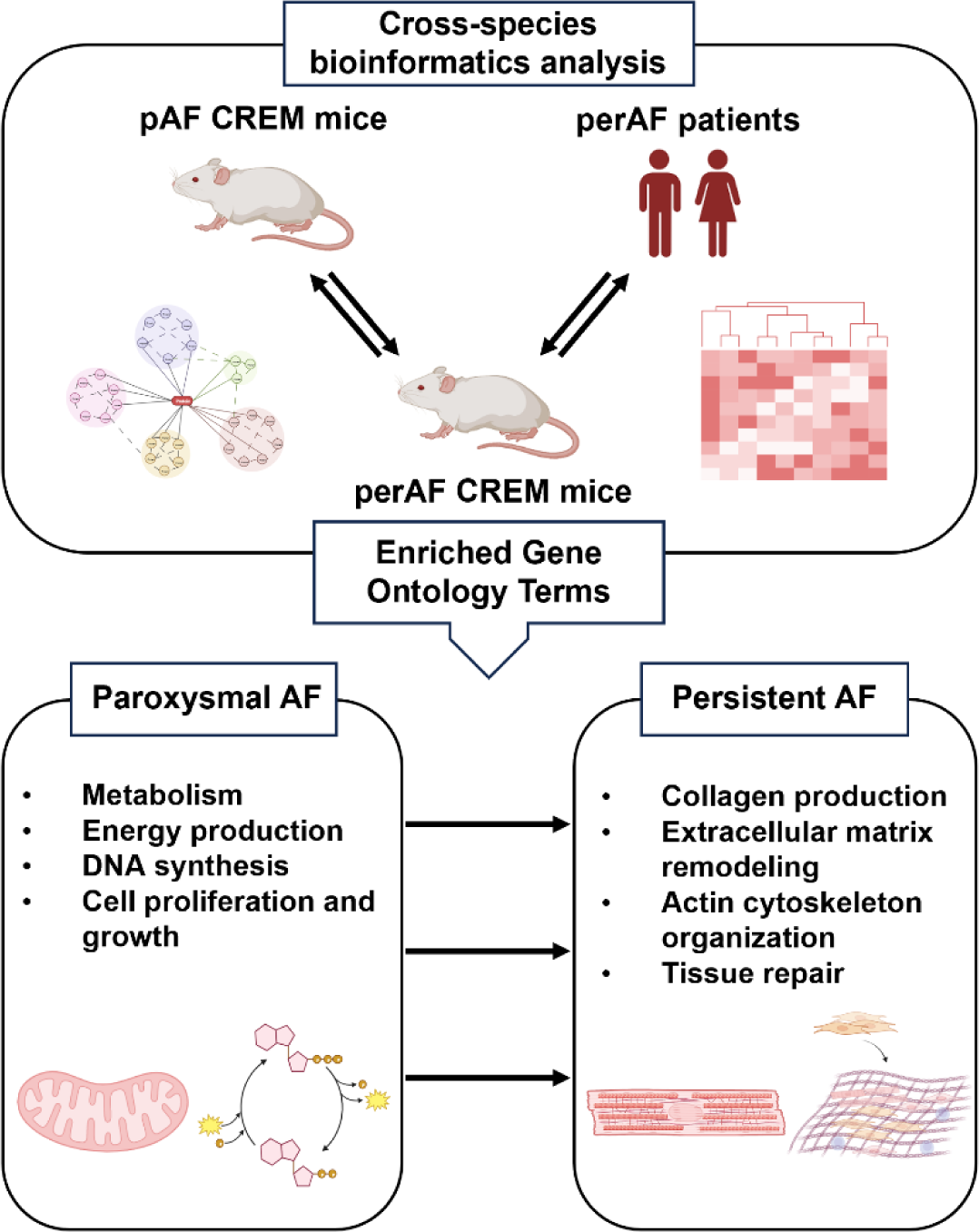

## 1. INTRODUCTION

Atrial fibrillation (AF) is the most common cardiac arrhythmia and a major risk factor for ischemic stroke and heart failure. According to data from the Global Burden of Disease study 2019, AF caused about 300,000 deaths while 59 million patients suffer from AF.[1] The progressive nature of disease progression complicates the assessment and treatment of AF.[2] AF typically begins as paroxysmal AF, characterized by intermittent arrhythmia periods lasting less than 7 days, and tends to progress over time to persistent AF (perAF). AF progresses over time and with age and more than half of all AF patients are >75 years old. [3]

AF is hallmarked by electrical and structural (fibrotic) remodeling in the myocardium, which underlies the progressive nature of the arrhythmia itself. AF related comorbidities, associated with structural changes in the myocardium, pose a particular challenge in AF treatment, as current therapies are largely focused on controlling electrical remodeling (the heart rate and rhythm), with no effective therapies targeting structural changes. Thus, further mechanistic understanding of how atrial (electrical and structural) remodeling contributes to the disease progression is necessary to develop more effective therapies for AF.[4] Electrical remodeling leading to ectopic firing and re-entrant activity of action potentials is considered to be causative for the onset of AF[5] as well as the maintenance of initial disease state. While it is shown that further structural and fibrotic remodeling is necessary for advanced disease state progression,[6] the underlying mechanisms are not well understood.

The use of animal models has enabled the study of specific genetic and molecular factors contributing to the pathogenesis of AF.[7, 8] While large animal models can recapitulate the disease progression seen in humans, they have some disadvantages like a longer time-course to develop advanced disease stages and difficulty to perform genetic modifications.[9] On the other hand, mouse models offer multiple advantages such as a shorter time course of disease development, ease of genetic manipulation, less genetic heterogeneity among animals, and overall cost-effectiveness. Several mouse models have been shown to recapitulate key disease features also seen in patients with AF.[7] For example, CREM transgenic (CREM-Tg) mice exhibit aberrant sarcoplasmic reticulum Ca^2+^ handling, which was demonstrated to serve as a mechanistic driver of AF progression.[10] CREM-Tg mice overexpress the CREM-IbΔC-X isoform, the expression of which is also elevated in human patients with perAF.[10] In these mice, atrial remodeling typically starts at 5-6 weeks of age and ectopic beats are observed at around 3 months, which is associated with spontaneous, paroxysmal AF.[8, 10, 11] As the CREM-Tg mice age, they develop atrial dilation, conduction abnormalities and extensive atrial fibrosis, which is associated with increasing persistence and burden of AF.[8, 10] The disease progression in CREM-Tg mice thus mimics the atrial remodeling and disease evolution often observed in patients with AF.

To gain a better understanding of the atrial protein landscape involved in the advanced stages of perAF, we performed unbiased mass spectrometry-based quantitative proteomics on 9-month-old CREM-Tg mice. Enrichment and network analysis identified ∼3,000 unique proteins in the atria of CREM-Tg mice and their age-sex matched wild-type littermate controls. Using multimodal analysis, the differentially expressed proteins (DEPs) were compared with publicly available datasets from young CREM-Tg mice.[12] While the young CREM-Tg mice with early-stage AF mainly exhibited changes in biological pathways involved in regulating metabolism, contractility, and electric activity in the atria, older CREM-Tg mice with more advanced-stage AF showed a significant shift in the gene expression patterns associated with fibrotic remodeling. The enrichment of actin filament assembly and ECM remodeling events emerged as a possible driver of alterations in atrial tissue integrity and contractility. Taken together, our study uncovered a novel shift in the atrial proteome associated with biological processes promoting atrial remodeling and fibrosis during more advanced stages of AF, similar to those observed in atrial tissue from patients with persistent AF. These pathways may be suitable for future studies with the goal of developing new therapeutic targets for AF prevention.

## 2. MATERIALS AND METHODS

A detailed description of all methods is provided in the **Supplemental Materials.** The authors did not use generative AI or AI-assisted technologies in the development of this manuscript.

### 2.1. Sample preparation for mass spectrometry

All animal studies were performed according to protocols approved by the Institutional Animal Care and Use Committee of Baylor College of Medicine conforming to the Guide for the Care and Use of Laboratory Animals published by the U.S. National Institutes of Health (NIH Publication No. 85-23, revised 1996). Mice with cardiomyocyte-specific overexpression of CREM-IbΔC-X (Tg) were described previously.[13] Mice were 9 months-of-age and divided into 2 experimental groups: 1) CREM-Tg mice and, 2) WT littermates. Mice were euthanized, and atria tissue was harvested, immediately snap frozen in liquid nitrogen, and stored at ™80 °C for downstream analysis. Tissue homogenization, protein digestion, peptide clean-up, and unbiased mass spectrometry were performed as detailed in the **Supplemental Material.**

### 2.2. Mass spectrometry (MS) data processing

The MS raw data was searched with Proteome Discoverer 2.1 software (Thermo Scientific, Waltham, MA) with Mascot 2.4 (Matrix Science, Chicago, IL) against NCBI refseq *Mus musculus* database (updated 03/24/2020). The following parameters were set for identification and quantification: 1) trypsin digestion with a maximum of two miscuts; 2) dynamic modification of oxidation (M), protein N-terminal acetylation, deamidation (NQ); 3) fixed modification of carbamidomethylation (C); 4) precursor mass tolerance of 20 ppm and fragment mass tolerance of 0.02 Da. The PSMs output file from Proteome Discoverer was grouped at the gene level. The gene product inference and iBAQ-based quantification was carried out using the gpGrouper algorithm to calculate peptide peak area (MS1) based expression estimates.[14] The median normalized and log10 transformed iBAQ values were used for data analysis.

### 2.3. Analysis of differentially expressed proteins

The differentially expressed proteins (DEPs) between the CREM-Tg and WT littermates were calculated using the moderated t-test to calculate p values and log2 fold changes in the R package “limma”.[15] The Benjamini-Hochberg method was used to adjust original p values. DEPs with an adjusted p value <0.05 and log2FC >1 or log2FC <-1 were considered as significant. The volcano plot was used to display DEPs. Principal component analysis (PCA) and unsupervised hierarchical clustering of distances between samples were performed to analyze sample clustering based on the protein expression profile similarities. PCA plot was generated by R package “ggplot2”[16] and R-package “pheatmap”[17] was used to generate hierarchically clustered heatmap. The common significant DEPs between our dataset and public datasets were obtained by the online Venn diagram tool (https://bioinformatics.psb.ugent.be/webtools/Venn/).

### 2.4. Function and pathway enrichment analysis of DEPs

DEP function and pathway enrichment analysis was carried out using the R package “clusterProfiler”.[18] Gene Ontology (GO) analysis was performed using the significant DEPs while the full list of DEPs was applied to Gene Set Enrichment Analysis. The Benjamini-Hochberg method was used to correct the p value and adjusted p <0.05 were considered statistically significant. Specifically, our GO analysis focused on the biological process (BP) category, which is a key aspect of protein function.

### 2.5. Protein-protein interaction network construction and module analysis

STRING database (https://string-db.org) was used to construct the protein-protein interaction (PPI) network of DEPs using the medium confidence score of 0.4 as a cut-off.[19] Cytoscape (version 3.9.1, http://www.cytoscape.org) was used for the network visualization.[20] Highly interconnected modules were extracted from the PPI network using the Cytoscape plug-in MCODE with the following criteria: degree cutoff = 2, node score cutoff = 0.2, k-core = 2, max depth = 100.[21]

### 2.6. Protein co-expression correlation analysis

Co-expression patterns between ECM proteins were determined based on the proteomics level. Pearson correlations coefficient between proteins were calculated using the R package “corrplot”.[22]

### 2.7. Identification of core proteins

Core proteins demonstrate complex interplays with other proteins in the processes. To prevent the selection bias, core proteins are defined based on their correlation relationship as well as their degrees of connection in the PPI network. The Deeply Integrated Human Single-Cell Omics (DISCO) database.[23] (https://www.immunesinglecell.org/) was used to verify the expression level of the core proteins in heart.

### 2.7. Dataset acquisition

The dataset from Seidl *et al.*[12] was obtained from 7-week-old mice with cardiomyocyte-specific overexpression of CREM-IbΔC-X (CREM-Tg). As controls, age-matched WT littermates from the same breeding colony were used. The dataset from Liu *et al*.[24] contains 18 human left atrial appendage (LAA) tissue samples including 9 with persistent AF and 9 with sinus rhythm. Different excel files containing differentially expressed proteins identified from each dataset were obtained.

### 2.8. Immunoblotting

Protein lysates were denatured for 10 minutes at 70°C in Laemmli buffer with beta-mercaptoethanol prior to electrophoresis on a 10% acrylamide gel. Proteins were transferred for 90 minutes at room temperature (RT) onto a PVDF membrane. Membranes were blocked 60 minutes at RT in OneBlock™ blocking buffer (#20-314, Genesee Scientific, Morrisville, NC) followed by overnight incubation at 4°C in primary antibodies for ITGAV (1:1,000, A2091, Abclonal, Woburn, MA), FBLN5 (1:1,000, A9961, Abclonal, Woburn, MA), LCP1 (1:1,000, A5561, Abclonal, Woburn, MA), and GAPDH (1:10,000 EMD Millipore, Burlington, MA) in OneBlock™ blocking buffer. After washing in TBS-Tween, membranes were incubated in goat anti-mouse IgG (H+L) superclonal™ secondary antibody, Alexa Fluor 680 (1:10,000, A28183, Thermo Fischer, Waltham, MA) or goat anti-rabbit IgG (H&L) antibody DyLightTM 800 conjugated (1:10,000, 611-145-002, Rockland, Limerick, PA) at a dilution of 1:10,000 in OneBlock™ blocking buffer prior to imaging with Li-Cor Odyssey Blot Imager.

### 2.9. Statistical analysis

Data were analyzed using analysis of variance (ANOVA) followed by the post hoc Bonferroni t-test for multiple group non-repeated measures. Independent groups in the same experiment were analyzed using unpaired *t*-test. Values are expressed as mean ± SEM, and p < 0.05 was considered statistically significant. All the statistical analyses were performed using GraphPad Prism 10 (GraphPad Software Inc., San Diego, CA) unless otherwise stated.

## RESULTS

### 3.1. Characterization of atrial dysfunction and remodeling in CREM-Tg mice

To examine the effects of cardiomyocyte-specific overexpression of CREM-IbΔC-X on atrial remodeling, echocardiographic studies were performed to assess atrial and ventricular function and structure. Representative long-axis echocardiography revealed clear images of the aortic root (Ao) and left atrium (LA) in 9-month-old CREM-Tg mice and WT littermates (**Fig. 1A**). The LA size was significantly larger in CREM-Tg mice compared to WT littermates (**Fig. 1B**). The atrial contractile function was studied using echo Doppler. Representative color Doppler images of the early and late flow peaks through the mitral valve showed an increased mitral early to after waves (E/A) ratio in CREM-Tg versus WT littermates (**Fig. 1C-D**), indicating abnormal ventricular filling probably because of decreased atrial contractility. The CREM-Tg mice also exhibited mild ventricular contractile dysfunction: the ejection fraction was reduced in CREM-Tg mice (51.4 ± 0.97) compared with WT littermates (61.7 ± 0.84; P=0.010). There were signs of left ventricular dilatation without increases in wall thickness, consistent with a mild dilated cardiomyopathy (**Supplemental Table 1**), consistent with prior studies. [25, 26] Furthermore, the amount of atrial fibrosis was examined using picrosirius red staining of longitudinal cardiac sections (**Fig. 1E**). Quantification revealed an increased amount of fibrosis in the atria from CREM-Tg compared with WT littermates (p = 0.006; **Fig. 1F**), consistent with prior studies [25].

**Figure 1.**
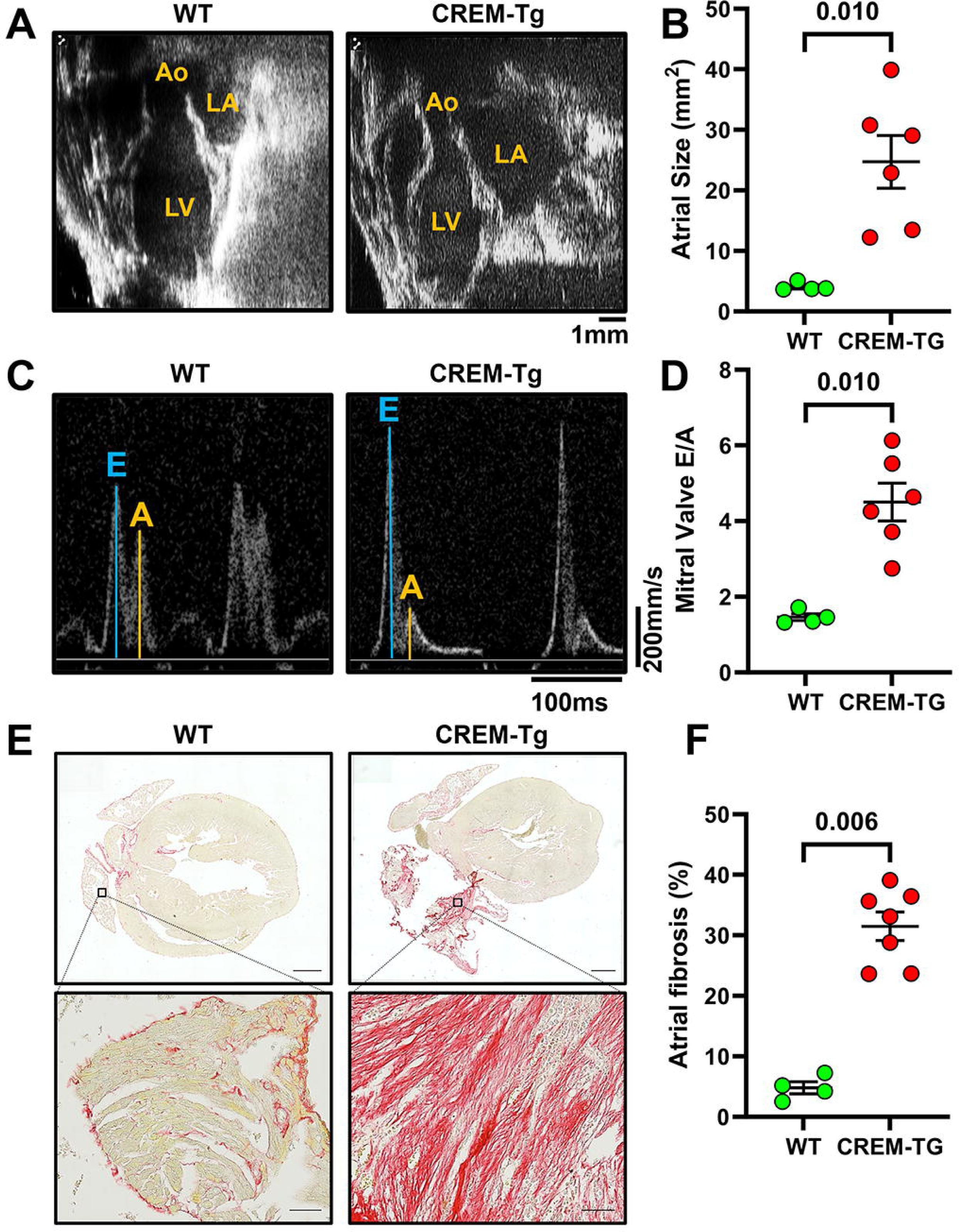
Atrial enlargement and fibrosis in CREM-Tg mice compared to WT littermates: **A**) Representative echocardiograph images and **B**) scatter plots showing increased atrial size in CREM-Tg mice (n=6) versus WT littermates (n=4). **C**) Echo Doppler recordings showing mitral valve flow and **D**) scatter plots showing increased mitral valve E/A ratios in CREM-Tg mice (n=6) versus WT littermates (n=4). **E**) Representative images of picrosirius red staining and **F**) quantification of interstitial fibrosis of atrial sections from CREM-Tg (n=7) versus WT control mice (n=4). Scale bar = 200μm (5X) and 50μm (20X). Data expressed as mean ± SEM. P values were determined using an unpaired *t* test.

### 3.2. Proteomic profiling and DEPs identification

To generate a complete profile of protein expression changes associated with chronic AF in CREM-Tg mice, we performed an unbiased label-free proteomic analysis. **Fig. 2** shows the workflow of this study, starting from sample preparation to mass spectrometry to eventual bioinformatic analysis of the results. To elucidate proteome alterations in mice with perAF, we analyzed atrial samples from three CREM-Tg mice and three WT littermates. Liquid chromatography with tandem mass spectrometry (LC-MS/MS) identified a total of 3,356 proteins. For the sake of reproducibility, the data set was filtered for detection of protein in all three replicates of either WT or CREM-Tg mice. Consequently, a total of 2,438 proteins were utilized for differential analysis.

**Figure 2.**
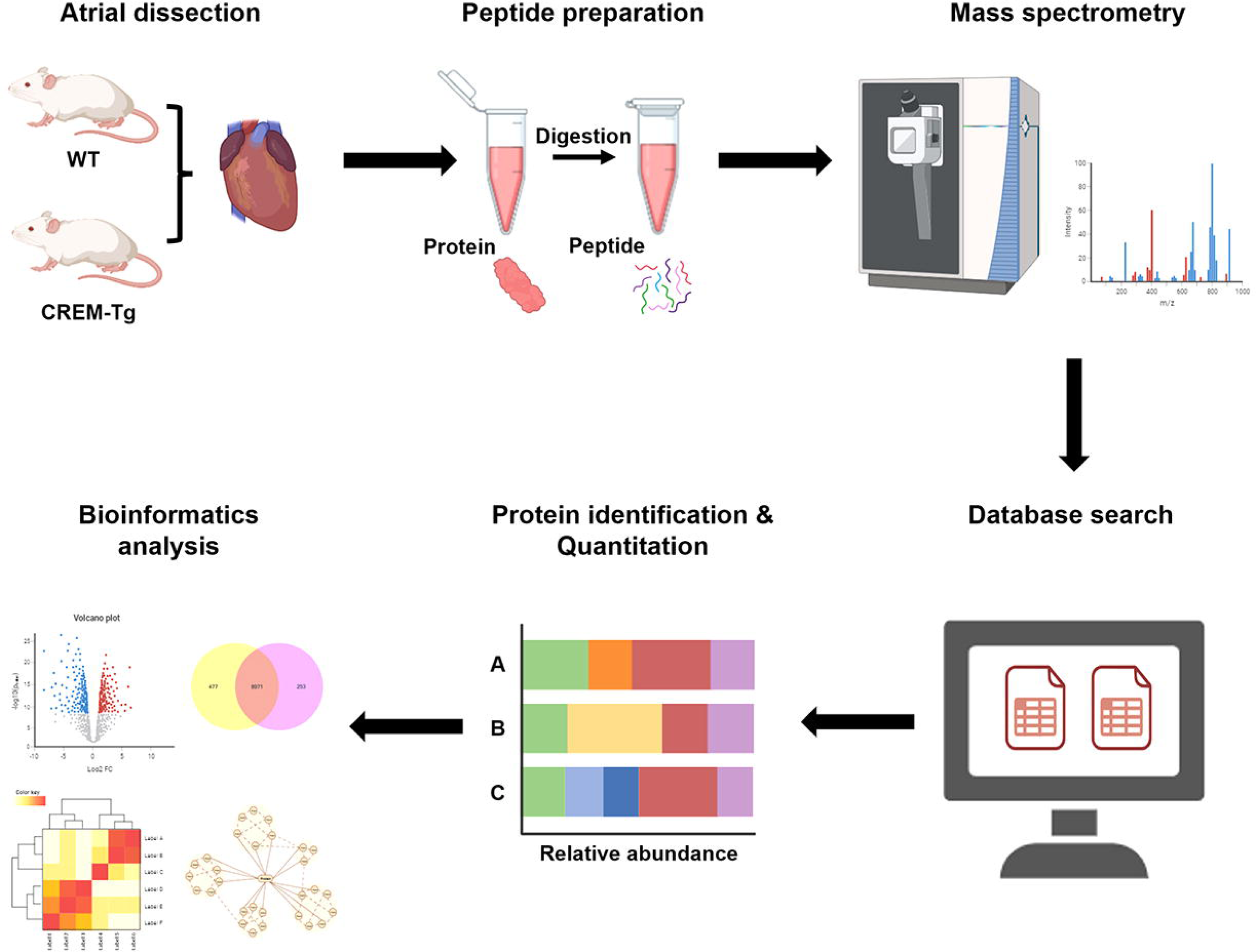
Workflow of the LC-MS/MS-based proteomic profiling of the atria from CREM-Tg mice with chronic AF and WT littermate controls in sinus rhythm.

To gain insight into the overall proteome alterations in CREM-Tg mice vs WT littermates, principal component analysis (PCA) and unsupervised clustering analysis of the overall protein expression levels identified by mass-spectrometry were performed (**Fig. 3A and Supplemental Fig. 1**). In the PCA plot, PC1 accounted for 89.1% of the variance in protein expression between samples while PC2 accounted for 9.3% of the variance, indicating differential separation between groups. Notably, the differential expression pattern of mouse CREM1 may be indicative of biological variability within the CREM-Tg group. To show the distribution of differentially expressed proteins (DEPs) between CREM- Tg mice and WT littermates in relation to their relative importance, a volcano plot was generated (**Fig. 3B**). Out of all 2,438 DEPs identified, 98 were significant using a cut-off filter of an adjusted p value <0.05 and log2FC >1 or <-1, respectively. Out of these, 82 significant DEPs were up-regulated, while 16 were down-regulated (**Supplemental Table 2**). Unsupervised clustering analysis revealed that the significant DEPs accurately separated the two group samples into two clusters, reinforcing the reliability of our differential expression analysis results (**Fig. 3C**). Subsequently, we selected all significant DEPs for further validation and downstream characterization.

**Figure 3.**
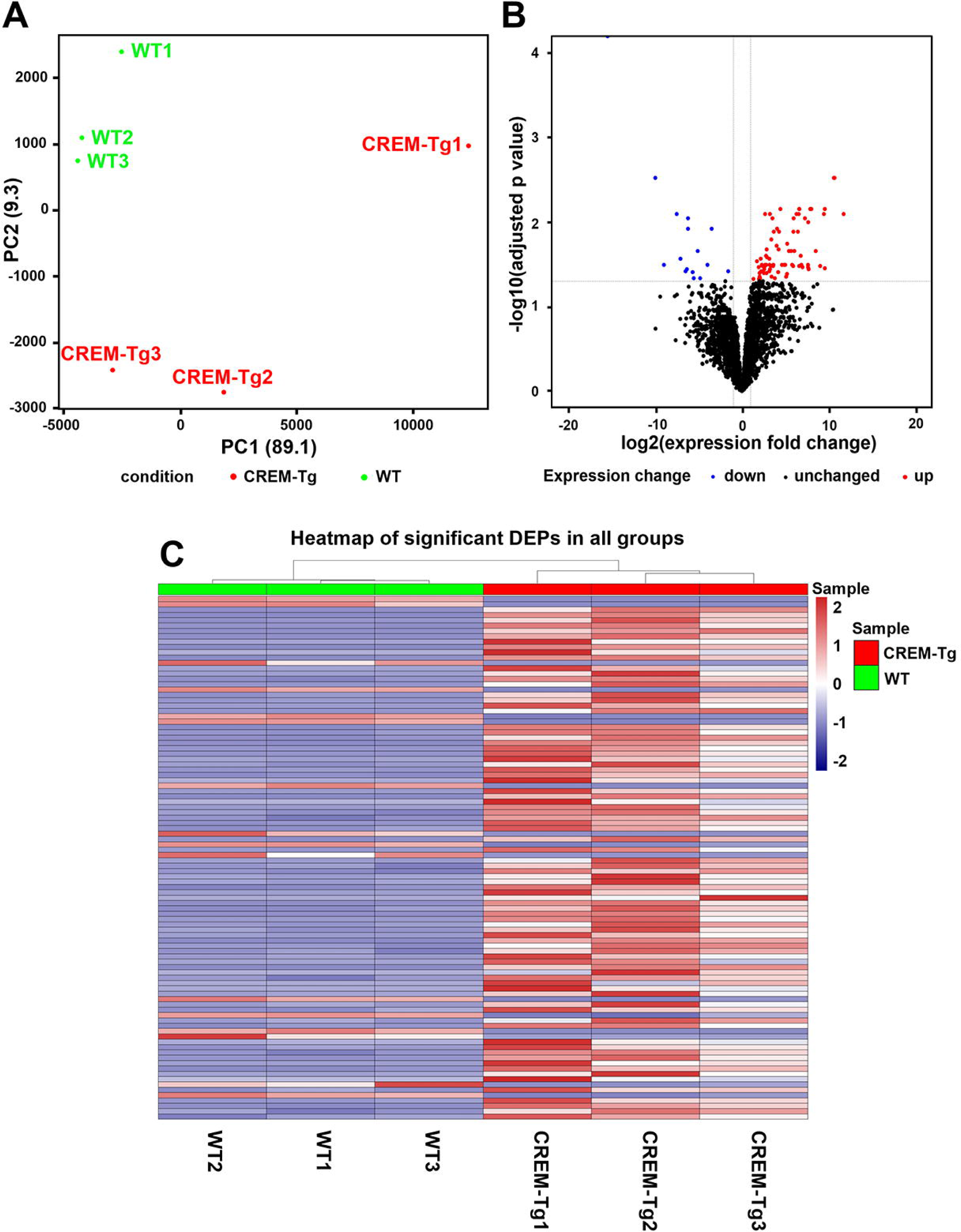
Multi-dimensional analysis of protein expression profile in CREM-Tg mice and WT littermates. **A**) Principal component analysis showing the overall effect of variances between the CREM-Tg mice and WT littermate controls. **B**) Volcano plot demonstrating the distribution of protein expression changes comparing CREM-Tg mice to WT littermates. Red dots represent significantly up-regulated proteins with adjusted p values <0.05 and log_2_FC >1 while blue dots represent significantly down-regulated proteins with adjusted p values <0.05 and log_2_FC <-1. **C**) Heatmap of differentially expressed proteins (DEPs) in each sample. The columns correspond to the samples and the rows correspond to individual proteins. Samples are grouped by clusters. The color scale is based on a *z*-score distribution from −2 (blue) to 2 (red), see scale. *Z*-scores were calculated on a protein-by-protein (row-by-row) basis by subtracting the mean and then dividing by the standard deviation.

### 3.3. Functional enrichment analysis of differentially expressed proteins

To identify biological functions that are involved in significant DEPs, we performed gene ontology (GO) enrichment analysis. We focused on the GO term ‘biological process’ (BP; GO:0008150) specifically. Significantly enriched BPs were classified into five clusters according to their functional descriptions and are shown in **Supplemental Fig. 2**. As shown in **Fig. 4A** our most enriched BPs were actin filament organization, extracellular matrix organization, extracellular structure organization, actin filament bundle assembly, etc., suggesting that the functions of significant DEPs are highly associated with actin dynamics and extracellular matrix (ECM) remodeling processes.

**Figure 4.**
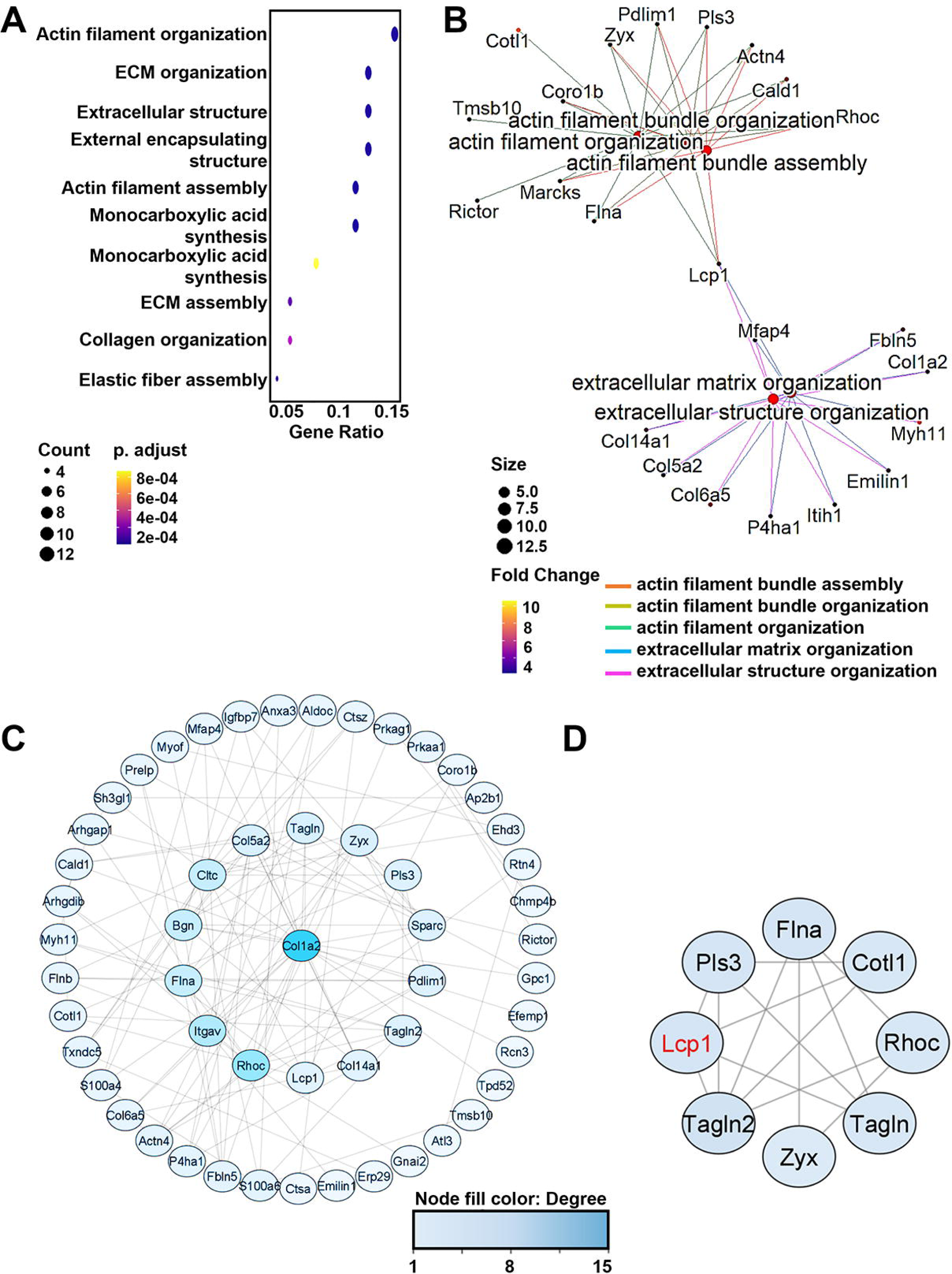
Pathway enrichment and PPI network analysis of differentially expressed proteins (DEPs). **A**) Dot plot showing the top 10 enriched biological processes (BPs) comparing CREM-Tg mice to WT littermates. **B**) The CNET plot depicting the proteins associated with the top 5 BPs. The log2 expression fold change overlaid as color gradient. **C**) PPI network of significant DEPs, in which the nodes represent proteins, and the edges represent the interaction between proteins. The darker the node, the higher the degree of the protein represented by the node within the network. **D**) Top 1 closely connected DEP module identified by the Molecular Complex Detection (MCODE) algorithm. Adjusted p values were calculated using the Benjamini-Hochberg procedure.

The gene concept network analysis revealed the association of proteins involved in the top 5 BP terms (**Fig. 4B**). The red nodes represent BP terms that are connected to DEPs. These DEPs were found to be enriched in relevant BPs. Furthermore, the color of each DEP node is indicative of their relative expression change. Notably, the protein LCP1 was found to be shared across all terms, signifying a promising protein for further investigation, and indicating that LCP1 may play a central role in regulating multiple diverse BPs involved in an advanced stage of AF. A full list of significantly enriched GO terms is available in **Supplemental Table 3**.

### 3.4. PPI network construction

A protein-protein interaction (PPI) network of all the significant DEPs was constructed to provide a systems-level perspective on protein-protein interactions and organization. The network contains 54 nodes and 118 interaction pairs (**Fig. 4C**). The top module within the network was identified through the application of Molecular Complex Detection (MCODE), which is a seed-and-extension approach. It initially assigns every node a weight, seeding these nodes to form initial clusters and expanding these clusters, eventually revealing functionally related clusters [21]. Notably, LCP1 was found in the top module (score = 4), highlighting its relevance as a candidate for further research (**Fig. 4D**).

### 3.5. Key proteins involved in atrial fibrotic remodeling

We recently identified an increased amount of interstitial fibrosis in the atria of 7-month-old CREM-Tg mice compared with WT littermates [25], which mimics the findings in patients with perAF. By controlling ECM remodeling and turnover, ECM dynamics play a crucial role in regulating fibrosis [11]. To further investigate mechanisms underlying ECM dynamics, their impact on AF, and to identify critical proteins involved in these processes, we next extracted DEPs from the enriched GO cluster annotated “collagen extracellular matrix structure”. In CREM-Tg mice, all DEPs in this cluster had elevated expression levels (**Fig. 5A**). The correlation analysis illustrated in **Fig. 5B** revealed a strong positive correlation between these ECM regulator proteins, indicating their expression may be regulated by shared factors and signaling pathways. We next constructed the PPI network of ECM DEPs. The network consists of 13 nodes and 28 interaction pairs (**Fig. 5C**). The degree of connection of each protein with other DEPs in the network is listed in the bar plot (**Fig. 5D**).

**Figure 5.**
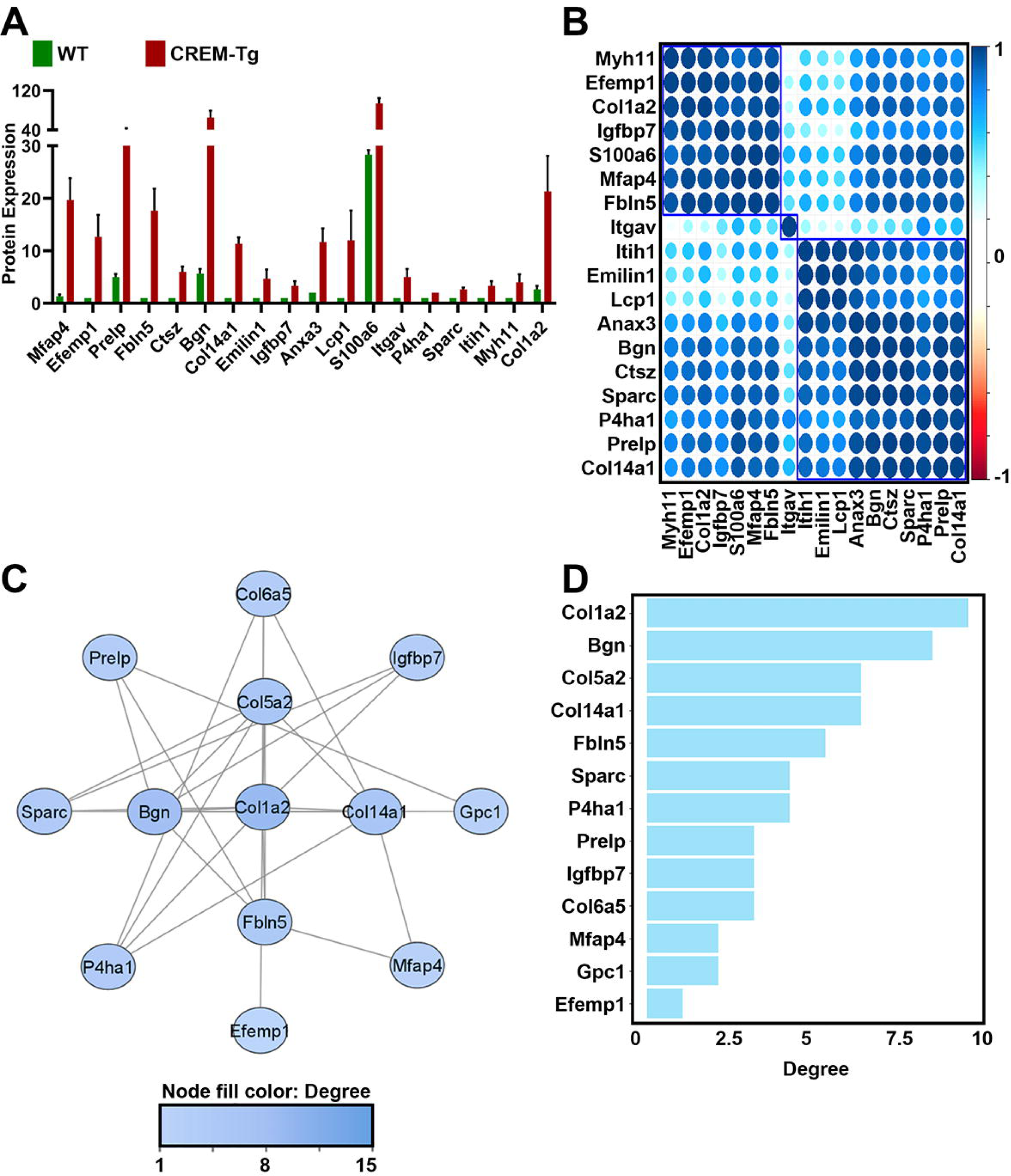
Protein co-expression and PPI network analysis of differentially expressed proteins in the extracellular matrix (ECM). **A**) Bar graphs showing normalized ECM protein expression levels of in the atrial of CREM-Tg mice and WT littermates. **B**) Dot plot shows the correlation analysis of ECM proteins. Only positive correlation between ECM proteins was observed. **C**) PPI network of ECM proteins. The darker the node, the higher the degree of the protein represented by the node in the network. **D**) Bar plot showing the degree value of each protein. Ordered from the largest to the smallest value. P values were determined using an unpaired *t* test.

### 3.6. Validation of key proteins

Considering the extensive characterization of collagen proteins in ECM remodeling and fibrosis [11, 27], we opted to focus our investigation on other proteins that may uncover new mechanistic pathways. Based on their abundance and the abovementioned analysis, we focused on ITGAV, LCP1, and FBLN5. We then referred to the DISCO database[23] for the expression of each key protein in curated single-cell RNA sequencing data from heart tissue samples. ITGAV and FBLN5 showed robust expression at the transcriptomic level within the heart tissue, primarily observed in the fibroblast subsets (**Supplemental Fig. 3 and 4**). While LCP1 expression could not be clearly observed in all the fibroblast subsets (**Supplemental Fig. 5**), considering its central location in our network analysis (**Fig. 4C**), we shortlisted these 3 proteins for further validation.

To validate our proteomics findings, we performed western blot analysis on ITGAV, FBLN5, and LCP1 expression in atrial tissue of CREM-Tg mice and WT littermates (**Fig. 6A,C,E**). We found that, consistent with the proteomics results, the protein expression levels of all three proteins were significantly higher in CREM-Tg mice versus WT littermates (ITGAV: 1.34±0.32; p=0.002; Lcp1: 0.75± 0.16; p<0.001; FBLN5: 0.49±0.16; p=0.010; **Fig. 6B,D,F**). These results lend support to the robustness of our quantitative proteomics analysis.

**Figure 6.**
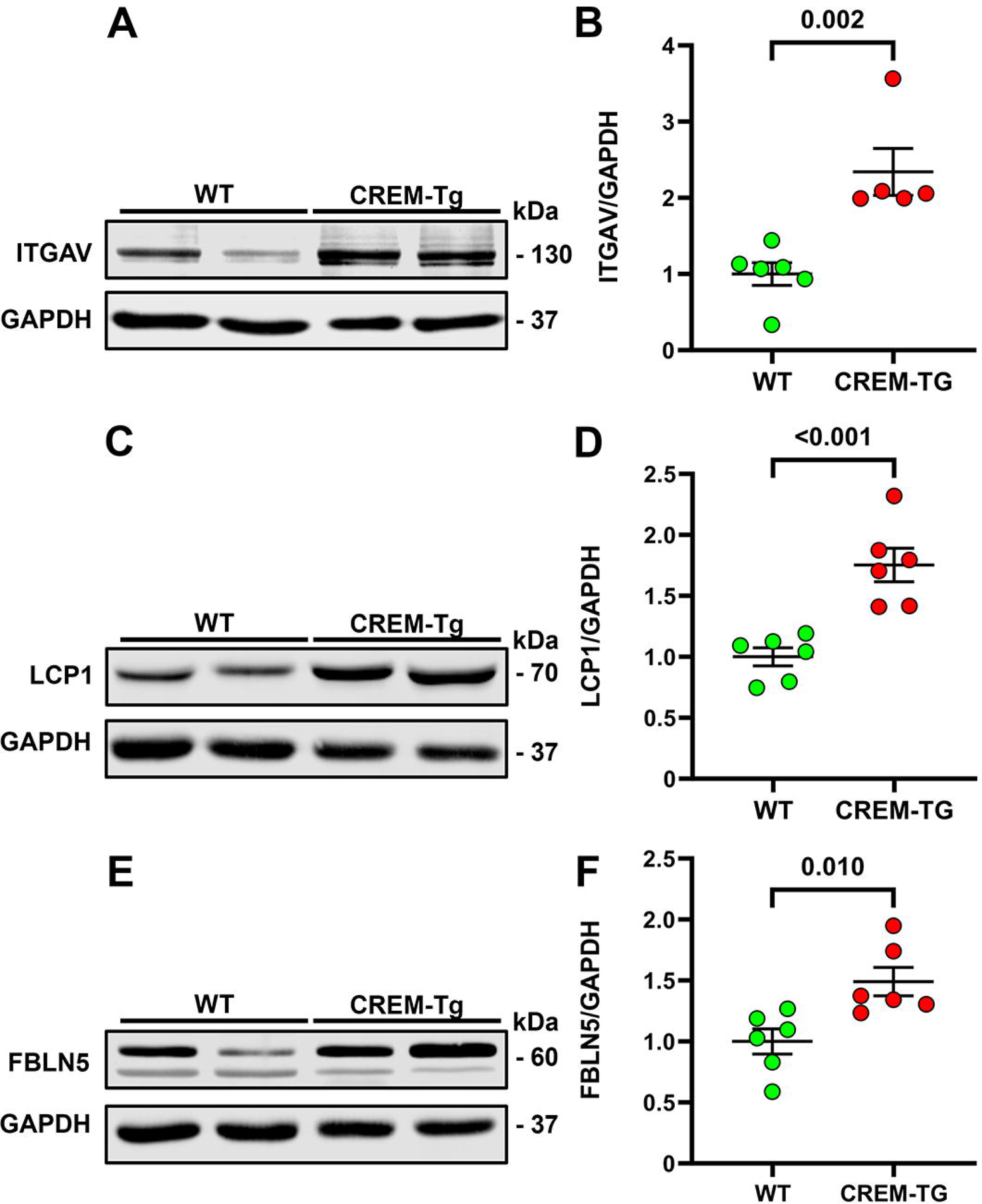
Western blot validation analysis of key ECM proteins in mouse atrial tissue. **A**) Western blot analysis of integrin alpha chain V (ITGAV) in atrial tissue of CREM-Tg mice and WT littermates. **B**) Quantification showing increased ITGAV levels in CREM-Tg mice. **C**) Western blot analysis of lymphocyte cytosolic protein 1 (LCP1) in atrial tissue and **D**) quantification showing increased LCP1 levels in CREM-Tg mice. **E**) Western blot analysis of fibulin-5 (FBLN5) in atrial tissue and **D**) quantification showing increased FBLN5 levels in CREM-Tg mice. Data expressed as mean ± SEM. P values were determined using an unpaired *t* test. Each dot represents an individual animal.

### 3.7. Comparison of DEPs from old vs young CREM-Tg mice

In a prior study, Seidl *et al.*[12] reported that biological processes enriched in young CREM-Tg mice at 7 weeks-of-age were mostly related to changes in metabolism, contractility, and electric activity. To assess whether there is a switch in the genetic program of CREM-Tg mice when AF becomes more persistent, we performed a comparative analysis between the datasets obtained from our CREM-Tg mice at 9 months-of-age and the publicly available datasets from Seidl *et al.*[12]. By overlapping DEPs, we found 14 to be common among the two datasets (**Fig. 7A**). Notably, all 14 of these DEPs showed a consistent pattern of expression change (**Fig. 7B**), i.e., 11 DEPs were overexpressed in both the datasets and 3 out of 14 were downregulated in both, with changes in the relative expression levels being more pronounced in the 9-month-old CREM-Tg mice. To understand the processes critical for regulating early vs. advanced stages of AF, we next conducted GO analysis on 94 DEPs that were unique to CREM-Tg mice at 7 weeks, 84 DEPs that were unique to Tg mice at 9 months, and 14 DEPs that were shared by both groups. While BPs related to metabolic regulation and energy production for normal muscle functioning were found to be unique for the 7-week-old CREM-Tg mice, we determined that as the AF progressed, there was a shift in the BPs to atrial remodeling and fibrosis (**Fig. 7C**). Interestingly, some of the BPs common for both early and late stages of AF were related to cytoskeletal organization and remodeling, suggesting that changes in the genes involved in processes related to atrial remodeling and ECM deposition begin in the early stages of AF, but are more pronounced as the disease progresses to advanced stages.

**Figure 7.**
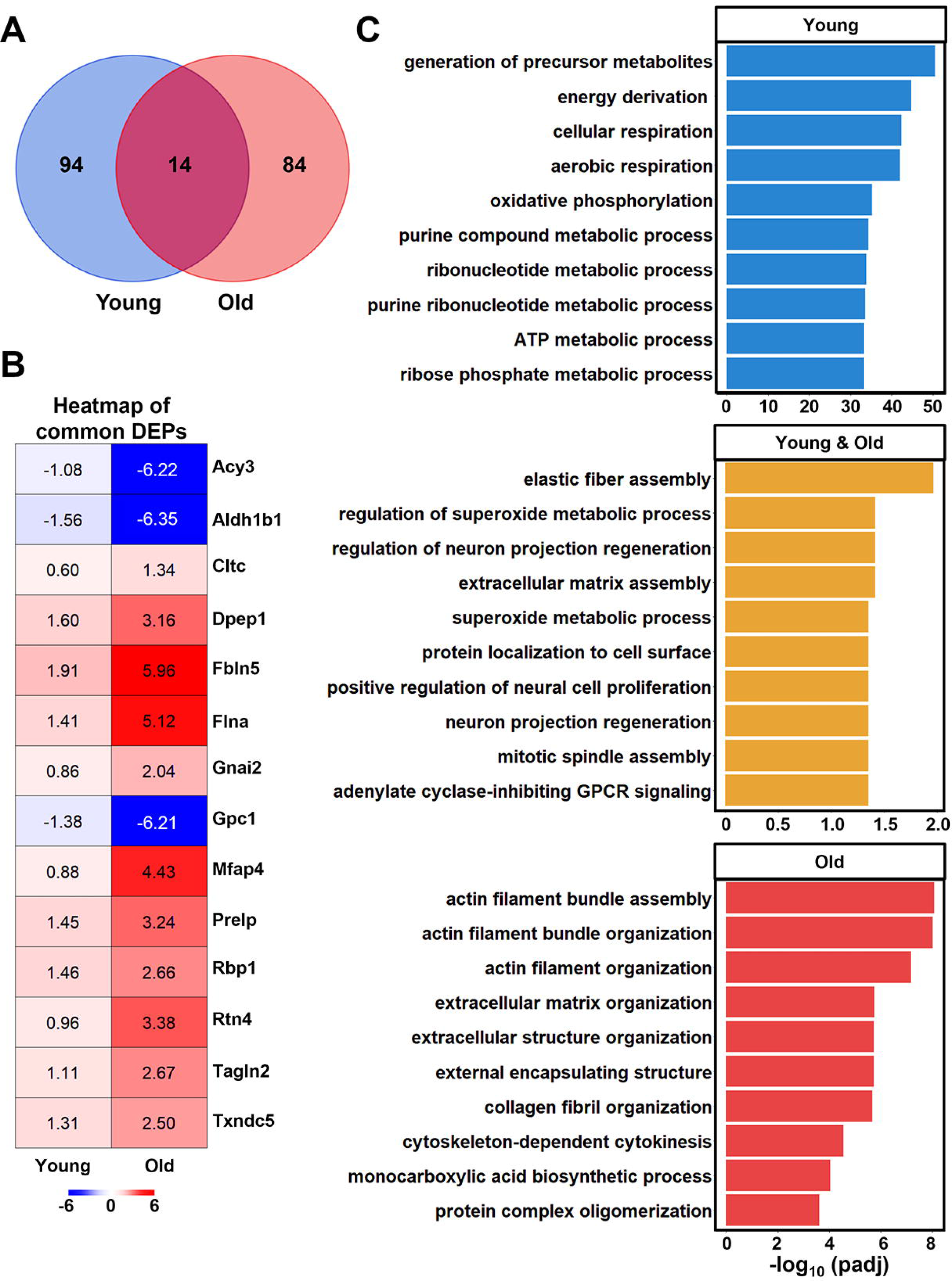
Differential expression profile (DEP) showing similarities and differences between young and old CREM-Tg mice. **A**) Venn diagram revealing common DEPs comparing old (9 months-of-age) and young (7 weeks-of-age) CREM-Tg mice, compared with WT littermates. **B**) Heatmap showing protein expression patterns in young and old CREM-Tg mice, compared with WT littermates. The color scale represents the log2 ratio of each protein in each group. **C**) Bar plots show the top 10 enriched biological processes using different subsets of DEPs. Blue color represents using DEPs only found in young CREM-Tg mice, red color represents DEPs only found in old CREM-Tg mice, and orange color represents DEPs found in both old and young CREM-Tg mice. Adjusted P values were calculated using the Benjamini-Hochberg procedure.

### 3.8. Comparison of DEPs from old CREM-Tg mice vs human AF patients

CREM-Tg mice have been reported as an effective AF model, demonstrating similarities in disease progression comparable to that observed in human AF.[11, 13, 25, 26, 28-30] To uncover additional areas of resemblance of molecular alterations in CREM-Tg mice in comparison with human patients, we obtained a publicly available dataset,[24] showing the atrial proteomic landscape in patients with persistent AF. We used the same significance threshold (fold change >1.2 and adjusted p value <0.1) as reported in this study and identified 480 significant DEPs in the human perAF patients compared to 98 in CREM-Tg mice (**Fig. 8**). Out of these, 21 DEPs were common across both the datasets (**Fig. 8A**). Remarkably, 18 of these 21 DEPs showed a consistent pattern of expression change (**Fig. 8B**), i.e., they were all overexpressed in both datasets. Many of those proteins (BGN, EMILIN1, IGFBP7, ITGAV, LCP1, SPARC) are involved in fibrogenesis processes, underscoring the relevance of CREM-Tg mice as a clinically relevant model to study persistent AF mechanisms.

**Figure 8.**
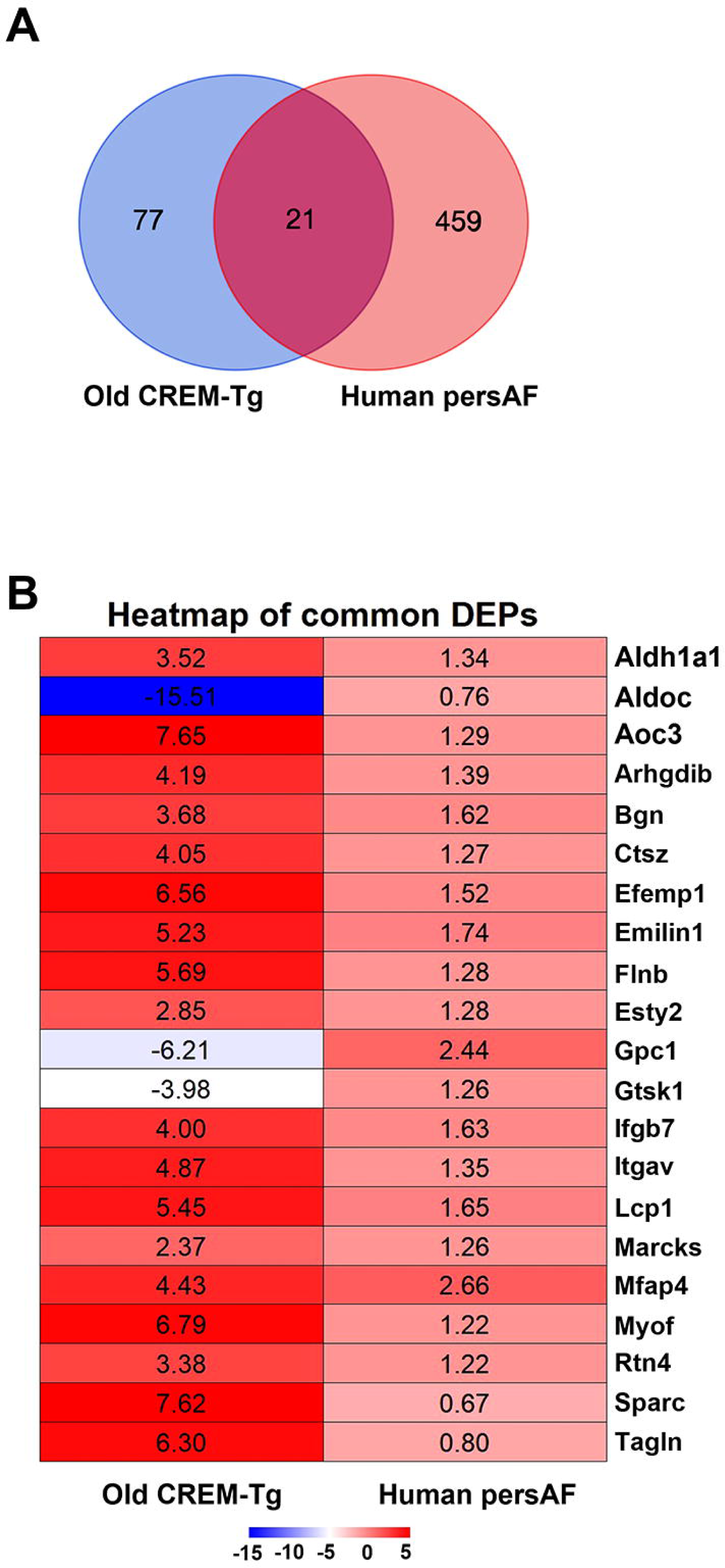
Protein expression changes in old CREM-Tg mice recapitulate those in human patients with persistent AF. **A**) Venn diagram reveals common differentially expressed proteins (DEPs) identified in old (9 months) CREM-Tg mice vs WT littermates, and patients with persistent atrial fibrillation (perAF) vs controls in sinus rhythm. B) Heatmap shows DEPs in old CREM-Tg mice and human patients with perAF. The color scale and the number in each box correspond to the log2 ratio of each protein in different groups. Adjusted P values were calculated using the Benjamini-Hochberg procedure.

## 4. DISCUSSION

In this study, we performed unbiased proteomic studies to uncover new molecular mechanisms that play a role in the pathogenesis of AF progression. Specifically, we utilized the well-characterized CREM-Tg mouse model of spontaneous AF development and progression that was previously shown to mimic major aspects of AF in humans, such as AF progression, atrial remodeling and fibrosis, and intracellular Ca^2+^ handling abnormalities.[11, 13, 25, 26, 28-30] Our results show that 9-month-old CREM-Tg mice with persistent AF develop enlarged atria and atrial fibrosis, mimicking the phenotype seen in patients with advanced persistent AF. Since the molecular pathways involved in atrial remodeling are not well understood, we performed mass spectrometry-based unbiased proteomic profiling of atrial tissues obtained from 9-month-old CREM-Tg mice. Unsupervised clustering analysis of differentially expressed proteins showed a clear distinction between CREM-Tg mice and their age-matched WT littermate controls. Gene ontology analysis of DEPs from CREM-Tg mice revealed upregulation of various proteins involved in actin-cytoskeleton organization as well as ECM deposition. A comparison with DEPs in the atria of young CREM-Tg mice showed that there was a clear switch from proteins involved in the regulation of metabolism and muscle contraction to fibrotic remodeling in mice with more advanced stages of AF. Moreover, comparative analysis with a publicly available proteomic dataset from patients with persistent AF showed an overlap with DEPs from CREM-Tg mice, and these overlapping genes were involved in biological processes related to atrial remodeling and fibrosis.

### 4.1. Enrichment of structural remodeling processes in AF

Enrichment analysis using DEPs from our proteomics results revealed that biological processes related to actin cytoskeleton organization and ECM remodeling were most significantly altered in CREM-Tg mice compared to the WT littermate controls, which is in concordance with clinical observations that advanced stage AF is associated with atrial structural remodeling including fibrosis, myocyte hypertrophy, and loss of contractile elements.[31] Excessive and disordered deposition of ECM creates complications such as fibrosis, which disrupts the normal architecture of the atrial myocardium, eventually creating a fibrotic substrate vulnerable to arrhythmic events.[32] Cytoskeletal proteins are thought to consist of the majority of proteins found in atrial cardiomyocytes.[33] Given the critical role cytoskeletal proteins play in regulating the contractile ability of cardiomyocytes,[34] the alteration in their expression will result in impaired cardiomyocyte contraction, prolonged structural damage, and disturbed atrial function. Changes in ECM remodeling and cytoskeleton organization significantly contribute to the structural remodeling events, compromising atrial function and electrical conduction. These alterations set the stage for the development and perpetuation of AF.

### 4.2. Role of actin-cytoskeleton organization in AF

The cytoskeleton of a cell is made of filamentous proteins such as actin, provides mechanical support to the cell, and is involved in various cellular processes, including stress transmission, cell shape change, and contractility.[35] Cytoskeletal rearrangement causes contractile dysfunction and impaired cellular function, suggesting the importance of cytoskeleton in maintaining normal cardiomyocyte function. The microtubule network, a key part of the cytoskeleton, is critical for maintaining balanced proteostasis in cardiomyocytes[36] and its derailment is shown to be involved in AF progression.[34, 37] Numerous studies have showed how cytoskeleton proteins contribute to the development of arrythmia, for example, alterations of cytoskeleton protein ankyrin B have been shown to cause ion channel dysfunction in heart, leading to increased susceptibility to arrythmias.[38-40] Mutations in actin-binding protein such as dystrophin disturbs voltage-dependent sarcolemmal ion channels, giving rise to arrhythmias.[41] In addition, reduced dystrophin protein was shown to associate with the reduced neuronal nitric oxide synthase, and increased AF inducibility.[42]

In our proteomics analysis, we revealed significant changes in the expression of proteins involved in the cytoskeleton dynamics in CREM-Tg mice compared to WT littermates, and some of the top regulated proteins include RhoC, MYH11, TAGLN, etc. Although there have been few investigations on the role of these proteins in AF, their potential as a promising target warrants more investigation. For example, dysregulation of RhoC signaling has been suggested to alter Ca^2+^ uptake of sarcoplasmic reticulum, potentially promoting AF pathogenesis in mice.[43] Additionally, a recent cross-ancestry meta-analysis involving over 1 million individuals discovered a previously unreported susceptibility locus in *MYH11* for AF, which highlights the possibility to delve into the role of MYH11 in AF pathogenesis.[44] Furthermore, in a prospective longitudinal cohort of over 4,000 patients, TAGLN, an actin-crosslinking protein, was identified as a putative causative protein for AF.[45] These findings emphasize the importance of focusing on these unique candidates in future functional studies of AF pathogenesis.

### 4.3. Role of ECM regulating proteins in AF

Atrial fibrosis has emerged as an important contributor to the pathophysiology of AF and has been associated with AF recurrences and the development of complications.[10, 46] The ECM primarily consists of fibrillar collagen including collagen types I and III, basement membrane proteins such as fibronectin, laminin, and fibrillin, and various proteoglycans.[47] Collagens are essential for maintaining the structural integrity of myocardial tissue, the basement membrane contributes to cell-cell interactions, while proteoglycans are crucial in adhesion and signaling processes. Our analysis revealed significant up-regulated expression of collagen proteins such as collagen I, VI, and XIV in CREM-Tg mice compared to WT littermates. The changes in the expression level of these collagens lead to their altered proportions in the ECM, which could contribute to the pathogenesis of atrial fibrosis, potentially leading to the development of AF.

In the present study, we validated the upregulation of ECM associated proteins – ITGAV, FBLN5, and LCP1 in CREM-Tg mice compared to WT littermates. ITGAV is known to play a central role in tissue fibrosis and its overexpression is thought to be the most upstream event of ECM deposition in fibroblasts.[48, 49] Pharmacological inhibition of ITGAV led to notable improvements in cardiac function in a mouse model of myocardial infarction.[50] Moreover, this inhibition resulted in a decreased expression of markers associated with cardiac fibroblast activation *in vitro*. Additionally, pro-fibrotic roles of ITGAV were reported in a hypertensive rat model.[51] FBLN5, an extracellular scaffold protein, is a known factor modulating deposition of elastin in ECM.[52] FBLN5 has been considered for its potential as an anti-fibrotic therapeutic target [53]. FBLN5 was identified as a novel protein highly expressed in patients with HBV/HCV-associated hepatic fibrosis in a large cohort study using quantitative proteomics.[54] Loss of FBLN5 resulted in reduced tissue stiffness and inflammation in a mouse model of cutaneous fibrosis.[53] suggesting that FBLN5 may be an ideal candidate to target profibrotic processes. Originally identified as an actin-binding protein in hematopoietic cells, [55] LCP1 has recently garnered attention for its involvement in tissue fibrosis. LCP1 was identified as a critical player in the liver fibrogenesis in non-alcoholic steatohepatitis through different integrative network-based analysis using publicly available bulk RNAseq datasets.[56] A recent study suggested that LCP1 contributes to pulmonary fibrosis by promoting NLRP3 inflammasome assembly in lung-resident alveolar macrophages.[57] Given the role of NLRP3 inflammasome in atrial fibrosis and AF,[30, 58] LCP1 may be a critical regulator of atrial fibrosis and remodeling.

Collectively, the involvement of these proteins highlights their crucial roles in regulating fibrosis, making them promising targets for anti-fibrotic therapeutic interventions. The roles of these proteins have never been reported in atrial fibrosis, underscoring the need for future investigations into their mechanisms and those of other DEPs found in our study in atrial fibrosis associated with AF. Understanding these mechanisms will expand our understanding of atrial fibrosis in AF and potentially identify novel therapeutic strategies for AF.

### 4.4. A shift in gene expression pattern from early to late-stage AF

We conducted a cross-species comparisons of atrial proteome data to deepen our understanding of the mechanisms driving AF initiation and progression. To identify possible age-dependent cellular alterations at the proteome expression level, we compared our dataset to a publicly available dataset of young CREM-Tg mice at 7 weeks of age.[12] A total of 14 common DEPs were identified, including several key players such as FBLN5, GPC1, MFAP4, PRELP, FLNA, and TAGLN2 involved in cytoskeleton organization or ECM dynamics. These proteins showed consistent changes in their regulation in two studies, indicating the onset of structural remodeling even at the early stage of AF, which was further exacerbated as AF progressed. GO analysis of these shared DEPs further revealed enrichment of processes closely related to ECM dynamics such as ECM assembly and elastic fiber assembly, whereas GO analysis of unique DEPs in young CREM-Tg mice showed they were primarily annotated to metabolic processes. This interesting finding suggests structural remodeling may occur at the molecular level prior to its observable manifestation. In line with our findings, Schulte *et al.*[59] reported more prominent atrial fibrosis and structural remodeling in 16-week-old CREM-Tg mice compared to younger 7-week-old CREM-Tg. Interestingly, their function and pathway enrichment analysis showed pathways related to actin cytoskeleton organization as well as critical processes in fibrotic progression such as cell adhesion and cell-matrix interaction enriched in younger CREM-Tg mice. The alignment of these findings reinforces the importance of structural remodeling in AF progression. Future interventions may include treating metabolic anomalies in the early stages of AF, while the observation that structural remodeling began early and persisted indicates its potential as a target throughout AF progression.

### 4.5. Clinical relevance of the experimental model of AF

Previous studies reported the CREM-Tg mouse as an effective model for studying AF [11, 25, 26, 29, 30]. However, these studies mainly focused on evaluating the suitability of the model through comprehensive biochemistry and physiological assessments. To further validate the clinical relevance of CREM-Tg mice, we conducted a comparative analysis between our dataset and that of a proteomics study using samples from human patients with persistent AF.[24] Our analysis revealed that 18 out of 21 shared DEPs showed consistent changes in expression, emphasizing the significance of these proteins. Many of these proteins were associated with the ECM remodeling, including BGN, ITGAV, EMILIN1, LCP1, SPARC, MFAP4, among others. These findings not only support the clinical relevance of our model but also highlight the potential for utility of these key proteins as biomarkers of AF or possible therapeutic targets. By targeting these proteins involved in ECM remodeling, it may be possible to stop the progression of AF and potentially improve patient outcomes.

### 4.6. Limitations and future directions

Our study points towards the potential of antifibrotic therapies in AF treatment. However, it is important to acknowledge limitations. For example, the number of samples included in the study is relatively small, in part due to the costs associated with these experiments; nevertheless, we still identified important molecular characteristics and establish the clinical significance of our experimental model of AF. Future functional studies need to be conducted to unravel mechanistic roles of these ECM proteins. Furthermore, in vivo fibroblast conditional knock-in or knock-out mouse model of these candidate proteins could provide further validation and strengthen our understanding of biological and structural changes in AF development and disease progression.

## CONCLUSION

In summary, we performed proteomics and bioinformatic analysis to demonstrate the involvement of the cytoskeleton and ECM remodeling in AF development We revealed the shifting landscape of the biological processes as the disease progresses and assessed the clinical significance of this experimental mouse model of AF. We identified candidate proteins that potentially regulate ECM remodeling and their association with AF disease progression. Our study paves the way for future novel strategies for the identification and prevention of AF.

## ABBREVIATIONS AND ACRONYMS

AF: atrial fibrillation
BP: biological process
CREM: cAMP-response element modulator DEP differentially expressed protein
ECG: electrocardiogram
ECM: extracellular matrix
FBLN5: fibulin 5
GO: gene ontology
GSEA: gene set enrichment analysis ITGAV integrin alpha V
LCP1: lymphocyte cytosolic protein 1
MS: mass spectrometry
pAF: paroxysmal atrial fibrillation
PCA: principal component analysis
perAF: persistent atrial fibrillation
PPI: protein-protein interaction
WT: wild type

## AUTHOR CONTRIBUTIONS

Study design: SZ, MH, XHTW; Execution/data collection: SZ, MH, YAS, SKL, EM, AJ; Data compilation and analysis: SZ, MH, YAS, SKL, AJ; Drafted manuscript: SZ and MH; Edited manuscript: all authors; Acquired funding, regulatory approvals: SR, XHTW.

## DATA AVAILABILITY

The mass spectrometry proteomics data have been deposited to the ProteomeXchange Consortium via the PRIDE partner repository with the dataset identifier PXD045882.

## APPENDIX A. SUPPLEMENTAL DATA

Supplemental data to this article can be found online at XXX.

