## Supplemental Materials for "Atrial Proteomic Profiling Reveals a Switch Towards Profibrotic Gene Expression Program in CREM-IbΔC-X Mice with Persistent Atrial Fibrillation"

[1]Cardiovascular Research Institute, Baylor College of Medicine, Houston, TX 77030, USA; [2]Department of Integrative Physiology, Baylor College of Medicine, Houston, TX 77030, USA; [3]Institute of Pharmacology and Toxicology, University of Münster, Münster, Germany; [4]Mass Spectrometry Proteomics Core, Baylor College of Medicine, Houston, TX, USA; [5]Department of Biochemistry, Baylor College of Medicine, Houston, TX 77030, USA; [6]Division of Cardiovascular Medicine, Radcliffe Department of Medicine, British Heart Foundation Centre of Research Excellence, NIHR Oxford BRC, University of Oxford, John Radcliffe Hospital, Oxford, UK; [7]Department of Medicine (in Cardiology), Baylor College of Medicine, Houston, TX 77030, USA; [8]Department of Neuroscience, Baylor College of Medicine, Houston, TX 77030, USA; [9]Department of Pediatrics (in Cardiology), Baylor College of Medicine, Houston, TX 77030, USA; [10]Center for Space Medicine, Baylor College of Medicine, Houston, USA.

### SUPPLEMENTAL METHODS

**Mass spectrometry analysis.** The cryo-pulverized tissue powder was lysed in 8M urea buffer. The protein lysate was reduced with 5mM dithiothreitol, alkylated with 10mM iodoacetamide and digested using LysC (#129-02541, Fujifilm Wako Chemicals, Richmond, VA) and trypsin (#90058, Thermo Fisher Scientific, San Jose, CA) enzyme mixture. The peptide desalting was carried out using Sep-Pak C18 columns (#WAT036945, Waters, Milford, MA) and quantified using the Pierce™ Quantitative Colorimetric Peptide Assay (#23275, Thermo Fisher Scientific, San Jose, CA). The peptides were dried in a speed vac and reconstituted in 5% methanol containing 0.1% formic acid buffer. The liquid chromatography with tandem mass spectrometry (LC-MS/MS) analysis was carried out using a nanoLC1200 system coupled to Orbitrap Exploris™ 480 (#BRE725539, Thermo Fisher Scientific, San Jose, CA) mass spectrometer. 1μg peptide was loaded on to a 1.9 μm Reprosil-Pur Basic C18 (#r13.b9, Dr.Maisch GmbH, Ammerbuch, Germany) 2 cm x 10 0 μm precolumn. The precolumn was switched in-line with an in-housed 20cm x 75 μ m analytical column packed with Reprosil-Pur Basic C18 equilibrated in 0.1% formic acid/water. The column heater was set to 55$^{o}$C. The peptide elution was done using a 110 min discontinuous gradient of 90% acetonitrile buffer (B) in 0.1% formic acid at 200nl/min (2-30%B: 86 min, 30-60%B: 6 min, 60-90%B: 9 min, 90-50%B: 9 min). The eluted peptides were directly electro-sprayed into the mass spectrometer. The full MS was acquired in Orbitrap (120000 resolution, scan range 350-1400 m/z, 3E6 normalized AGC target and 50 ms maximum injection time). The MS/MS for the top 50 dependent scan was done in Orbitrap (15000 resolution, 1.2 m/z isolation window, 1E5 normalized AGC target and 50 ms maximum injection time). The HCD fragmentation energy was 32%. The dynamic exclusion was set to 15 seconds.[1, 2]

**Echocardiography.** Mice were anesthetized using isoflurane (1.5%-2% v/v in 100% oxygen). After removal of the chest hair using Nair cream, mice were positioned onto a heated platform to maintain body temperature around 37.0 ± 0.5 °C.[3] Ultrasound echocardiography was conducted with the Visual Sonics Vevo F2 using a 57 MHz probe (FujiFilm VisualSonics, Toronto, ON, Canada). Both B-mode and M-mode images of the heart in short-axis orientation were captured to evaluate systolic function. To assess the area of the left atrium, we also captured long-axis B-mode images of the heart. The Vevo LAB software was used to analyze the short axis M-mode images to obtain systolic parameters. The long-axis B-mode images were further analyzed to measure the left atrial size. Next, we utilized color Doppler and pulsed wave Doppler in 4-chamber view to evaluate the flow through the mitral valve.[4, 5] The color Doppler images were analyzed to determine early (E) and subsequent (A) waves through mitral valve.

**Picrosirius red staining.** The heart tissues were fixed in 10% formalin solution (#HT501128-4L, Sigma, St Louis MO) for 48 hours and washed three times with PBS 1x, then added to 70% ETOH, and further processed in a tissue processor for paraffin embedding. The paraffin sections (7 µms) were deparaffinized by placing them into 3 xylene changes for 3 minutes each, followed by a gradient ETOH rehydration to DI water. For the picrosirius red Assay (#ab245887, Abcam Waltham, MA), the tissue samples were moved from DI water to Phosphomolybdic Acid Solution (0.2%) for 5 minutes, then the slides were placed in picrosirius red solution for 90 minutes, and finally added to 2 changes of Acetic Acid Solution. Afterwards, the slides were dehydrated in a gradient of ETOH and fixed in 3 changes of Xylene for 1 minute each. The tissue slides were sealed with a xylene based mounting media (#8312-4, Thermo-Fisher Waltham, MA) to protect from fading. Images were taken with ZEISS Axioscope 5 Smart Laboratory Microscope (Zeiss, Oberkochen, Germany), and analyzed using ImageJ (NIH, Bethesda, MD).

**Supplemental Table 1.** Echocardiographic parameters of chronic CREM-Tg and WT mice.

| | **WT (n = 4)** | **CREM-Tg (n = 6)** | **p value** |
|---|---|---|---|
| **Heart rate (bpm)** | 551.1 ± 27.1 | 508.9 ± 35.8 | 0.352 |
| **LVESD (mm)** | 2.79 ± 0.07 | 3.57 ± 0.06 | **0.010** |
| **LVEDD (mm)** | 4.16 ± 0.08 | 4.85 ± 0.07 | **0.010** |
| **LVESV (μL)** | 29.5 ± 1.81 | 53.6 ± 2.25 | **0.010** |
| **LVEDV (μL)** | 76.8 ± 3.42 | 110.2 ± 3.5 | **0.010** |
| **SV (μL)** | 47.3 ± 1.70 | 56.6 ± 1.83 | **0.010** |
| **LVEF (%)** | 61.7 ± 0.84 | 51.4 ± 0.97 | **0.010** |
| **LVFS (%)** | 32.8 ± 0.56 | 26.3 ± 0.59 | **0.010** |
| **CO (mL/min)** | 26.0 ± 0.87 | 28.7 ± 1.72 | 0.257 |
| **LVAW;s (mm)** | 0.89 ± 0.03 | 0.84 ± 0.03 | 0.352 |
| **LVAW;d (mm)** | 0.57 ± 0.03 | 0.59 ± 0.04 | 0.800 |
| **LVPW;s (mm)** | 0.95 ± 0.02 | 0.92 ± 0.05 | 0.610 |
| **LVPW;d (mm)** | 0.60 ± 0.01 | 0.59 ± 0.03 | 0.914 |
| **LA (mm2)** | 4.08 ± 0.35 | 24.7 ± 4.36 | **0.010** |
| **MV E/A ratio** | 1.46 ± 0.09 | 4.50 ± 0.51 | **0.010** |

Bpm, beats per minute; CO, cardiac output; LA, left atrial diameter; LVAW;d, left ventricular anterior wall diameter in diastole; LVAW;s, left ventricular anterior wall diameter in systole; LVEDD, left ventricular end-diastolic diameter; LVEDV, left ventricular end-diastolic volume; LVEF, left ventricular ejection fraction; LVESD, left ventricular end-systolic diameter; LVESV, left ventricular end-systolic volume; LVFS, left ventricular fractional shortening; LVPW;s, left ventricular posterior wall diameter in systole; LVPW;d, left ventricular posterior wall diameter in diastole; MV E/A ratio, ratio of E velocity to

A velocity; SV, stroke volume. Unpaired t test was performed to compare the two groups. Data expressed as mean ± SEM. Statistically significant P values ($P<0.05$) are shown in bold.

**Supplemental Table 2.** List of significant differentially expressed proteins.

| Protein | log2FoldChange | padj |
|---|---|---|
| **Aldoc** | -15.509 | <0.001 |
| **Ndufb2** | -10.002 | 0.003 |
| **Rhoc** | 10.628 | 0.003 |
| **Rcn3** | 10.695 | 0.003 |
| **Mfap4** | 4.433 | 0.007 |
| **Cald1** | 6.612 | 0.007 |
| **Prkag1** | 7.883 | 0.007 |
| **Aldh1a3** | 8.014 | 0.007 |
| **Ehd3** | 9.575 | 0.007 |
| **Tomm34** | 6.646 | 0.007 |
| **Orm1** | 11.734 | 0.008 |
| **Actn4** | 2.68 | 0.008 |
| **Camk1d** | -7.514 | 0.008 |
| **Atp1a3** | 9.458 | 0.008 |
| **Efemp1** | 6.558 | 0.008 |
| **Prelp** | 3.243 | 0.008 |
| **Tagln** | 6.297 | 0.008 |
| **Acy3** | -6.221 | 0.009 |
| **Aldh1a1** | 3.52 | 0.009 |
| **Fbln5** | 5.964 | 0.009 |
| **Prkaa1** | 7.273 | 0.009 |
| **Aoc3** | 7.65 | 0.010 |
| **Nploc4** | -3.479 | 0.012 |
| **Gpc1** | -6.214 | 0.012 |
| **Ctsz** | 4.052 | 0.012 |
| **Bgn** | 3.676 | 0.013 |
| **Pls3** | 4.281 | 0.013 |
| **Col14a1** | 5.955 | 0.013 |
| **Creg1** | 6.459 | 0.013 |
| **Erp29** | 3.401 | 0.016 |
| **Emilin1** | 5.233 | 0.018 |
| **Igfbp7** | 3.999 | 0.019 |
| **Anxa3** | 2.874 | 0.021 |
| **Erlin2** | 4.247 | 0.021 |
| **Lcp1** | 5.446 | 0.022 |
| **Hsd17b13** | -5.099 | 0.022 |
| **Gimap4** | 6.04 | 0.022 |
| **Esyt2** | 5.692 | 0.022 |
| **Cotl1** | 8.515 | 0.022 |
| **Myof** | 6.795 | 0.025 |
| **Septin2** | 2.792 | 0.025 |
| **Pdlim1** | 2.881 | 0.026 |

| | | |
|---|---|---|
| **Zyx** | 2.166 | 0.027 |
| **Tmsb10** | 3.161 | 0.027 |
| **Dido1** | -7.092 | 0.027 |
| **S100a6** | 1.729 | 0.029 |
| **Gstk1** | -3.976 | 0.032 |
| **Tagln2** | 2.674 | 0.032 |
| **Sds** | -8.993 | 0.032 |
| **Fmo2** | 3.198 | 0.032 |
| **Aldh1a7** | 2.823 | 0.032 |
| **Chmp4b** | 2.311 | 0.032 |
| **LOC100862446** | 3.633 | 0.032 |
| **Ftl1** | 3.633 | 0.032 |
| **Psmd12** | 2.713 | 0.032 |
| **Atl3** | 5.799 | 0.032 |
| **Dnajc5** | 6.863 | 0.032 |
| **Itgav** | 4.872 | 0.032 |
| **P4ha1** | 4.62 | 0.032 |
| **Sparc** | 7.62 | 0.032 |
| **Nedd4l** | 5.009 | 0.032 |
| **Rbp1** | 2.659 | 0.033 |
| **Txndc5** | 2.502 | 0.033 |
| **Coro1b** | 2.596 | 0.033 |
| **Selenof** | 5.92 | 0.033 |
| **Marcks** | 2.367 | 0.033 |
| **Rtn4** | 3.384 | 0.033 |
| **Itih1** | 3.61 | 0.033 |
| **Col6a5** | 6.606 | 0.033 |
| **Hmgn1** | 9.034 | 0.033 |
| **Sh3gl1** | 6.995 | 0.034 |
| **Septin7** | 1.949 | 0.034 |
| **Dad1** | 7.624 | 0.034 |
| **S100a4** | 9.581 | 0.035 |
| **Myh11** | 7.703 | 0.036 |
| **Aldh1b1** | -6.353 | 0.036 |
| **Col1a2** | 3.249 | 0.036 |
| **Arhgap1** | 2.589 | 0.036 |
| **Ly6g6f** | -1.579 | 0.038 |
| **Dpep1** | 3.157 | 0.038 |
| **Ctsa** | 2.222 | 0.038 |
| **Tex30** | -6.508 | 0.038 |
| **Armh1** | -5.675 | 0.039 |
| **Arhgdib** | 4.193 | 0.039 |
| **Gnai2** | 2.041 | 0.040 |
| **Samhd1** | 2.543 | 0.040 |
| **Flnb** | 2.846 | 0.040 |
| **Rictor** | 5.198 | 0.041 |
| **Tpd52** | 2.045 | 0.044 |

| | | |
|---|---|---|
| **Kctd12** | 3.428 | 0.044 |
| **Flna** | 5.117 | 0.044 |
| **Hddc3** | -4.827 | 0.046 |
| **Ptbp1** | 2.096 | 0.046 |
| **Mcee** | -5.552 | 0.046 |
| **Col5a2** | 3.804 | 0.046 |
| **Cltc** | 1.341 | 0.047 |
| **Ap2b1** | 1.981 | 0.047 |
| **C3** | 1.991 | 0.048 |

Proteins abundance was expressed as Log2 fold change > 1, or < -1 and adjusted P < 0.05 comparing between CREM-Tg and WT mice. Adjusted P values were calculated using the Benjamini-Hochberg procedure.

**Supplemental Table 3.** List of significantly enriched Gene Ontology (biological processes) terms.

| ID | Description | p.adj |
|---|---|---|
| **GO:0051017** | actin filament bundle assembly | 1.95E-07 |
| **GO:0061572** | actin filament bundle organization | 1.95E-07 |
| **GO:0007015** | actin filament organization | 1.41E-06 |
| **GO:0030198** | extracellular matrix organization | 2.02E-06 |
| **GO:0043062** | extracellular structure organization | 2.02E-06 |
| **GO:0045229** | external encapsulating structure organization | 2.02E-06 |
| **GO:0048251** | elastic fiber assembly | 1.23E-05 |
| **GO:0085029** | extracellular matrix assembly | 6.58E-05 |
| **GO:0030199** | collagen fibril organization | 0.001 |
| **GO:0072330** | monocarboxylic acid biosynthetic process | 0.001 |
| **GO:0002138** | retinoic acid biosynthetic process | 0.001 |
| **GO:0016102** | diterpenoid biosynthetic process | 0.001 |
| **GO:0016114** | terpenoid biosynthetic process | 0.004 |
| **GO:0046394** | carboxylic acid biosynthetic process | 0.005 |
| **GO:0016053** | organic acid biosynthetic process | 0.005 |
| **GO:0042573** | retinoic acid metabolic process | 0.011 |
| **GO:0022604** | regulation of cell morphogenesis | 0.011 |
| **GO:0006001** | fructose catabolic process | 0.011 |
| **GO:0051259** | protein complex oligomerization | 0.015 |
| **GO:0032956** | regulation of actin cytoskeleton organization | 0.015 |
| **GO:0008299** | isoprenoid biosynthetic process | 0.016 |
| **GO:0002072** | optic cup morphogenesis involved in camera-type eye development | 0.016 |
| **GO:0036438** | maintenance of lens transparency | 0.016 |
| **GO:1902396** | protein localization to bicellular tight junction | 0.016 |
| **GO:1902903** | regulation of supramolecular fiber organization | 0.017 |
| **GO:0110053** | regulation of actin filament organization | 0.019 |
| **GO:0061640** | cytoskeleton-dependent cytokinesis | 0.021 |
| **GO:0051260** | protein homooligomerization | 0.021 |
| **GO:1990748** | cellular detoxification | 0.021 |
| **GO:0032970** | regulation of actin filament-based process | 0.021 |
| **GO:0086036** | regulation of cardiac muscle cell membrane potential | 0.021 |
| **GO:0045765** | regulation of angiogenesis | 0.024 |
| **GO:0033572** | transferrin transport | 0.024 |
| **GO:0110095** | cellular detoxification of aldehyde | 0.024 |
| **GO:1901342** | regulation of vasculature development | 0.025 |
| **GO:0060900** | embryonic camera-type eye formation | 0.027 |
| **GO:0097237** | cellular response to toxic substance | 0.027 |
| **GO:0031345** | negative regulation of cell projection organization | 0.029 |
| **GO:0098693** | regulation of synaptic vesicle cycle | 0.031 |
| **GO:0032963** | collagen metabolic process | 0.031 |
| **GO:0045807** | positive regulation of endocytosis | 0.033 |

| | | |
|---|---|---|
| **GO:0001667** | ameboidal-type cell migration | 0.033 |
| **GO:0051639** | actin filament network formation | 0.033 |
| **GO:0060249** | anatomical structure homeostasis | 0.034 |
| **GO:0099504** | synaptic vesicle cycle | 0.034 |
| **GO:0098754** | detoxification | 0.034 |
| **GO:0051764** | actin crosslink formation | 0.035 |
| **GO:0030100** | regulation of endocytosis | 0.036 |
| **GO:0033690** | positive regulation of osteoblast proliferation | 0.038 |
| **GO:0110096** | cellular response to aldehyde | 0.038 |
| **GO:0007030** | Golgi organization | 0.038 |
| **GO:0031589** | cell-substrate adhesion | 0.041 |
| **GO:0006000** | fructose metabolic process | 0.041 |
| **GO:0019318** | hexose metabolic process | 0.043 |
| **GO:0010224** | response to UV-B | 0.043 |
| **GO:0048268** | clathrin coat assembly | 0.043 |
| **GO:0034976** | response to endoplasmic reticulum stress | 0.043 |
| **GO:0001523** | retinoid metabolic process | 0.043 |
| **GO:0090307** | mitotic spindle assembly | 0.043 |
| **GO:0060348** | bone development | 0.043 |
| **GO:0099003** | vesicle-mediated transport in synapse | 0.043 |
| **GO:0019216** | regulation of lipid metabolic process | 0.043 |
| **GO:0016052** | carbohydrate catabolic process | 0.043 |
| **GO:0046348** | amino sugar catabolic process | 0.043 |
| **GO:0016101** | diterpenoid metabolic process | 0.043 |
| **GO:1901136** | carbohydrate derivative catabolic process | 0.044 |
| **GO:0034394** | protein localization to cell surface | 0.044 |
| **GO:0005996** | monosaccharide metabolic process | 0.044 |
| **GO:0071786** | endoplasmic reticulum tubular network organization | 0.044 |
| **GO:0010977** | negative regulation of neuron projection development | 0.044 |
| **GO:0048488** | synaptic vesicle endocytosis | 0.044 |
| **GO:0140238** | presynaptic endocytosis | 0.044 |
| **GO:2000179** | positive regulation of neural precursor cell proliferation | 0.045 |
| **GO:0034764** | positive regulation of transmembrane transport | 0.045 |
| **GO:0051258** | protein polymerization | 0.045 |
| **GO:0000281** | mitotic cytokinesis | 0.046 |
| **GO:0006721** | terpenoid metabolic process | 0.048 |

Adjusted P values for comparisons between CREM-Tg and WT mice were calculated using the Benjamini-Hochberg procedure ($p < 0.05$ considered to be significant).

**Supplemental Figure 1. Heatmap of the expression profiles of all DEPs in each sample.** The columns correspond to the samples and the rows correspond to the proteins. Samples are grouped by clusters. The color scale is based on a z-score distribution from −2 (blue) to 2 (red). DEP: differentially expressed protein. Z-scores were calculated on a protein-by-protein (row-by-row) basis by subtracting the mean and then dividing by the standard deviation.

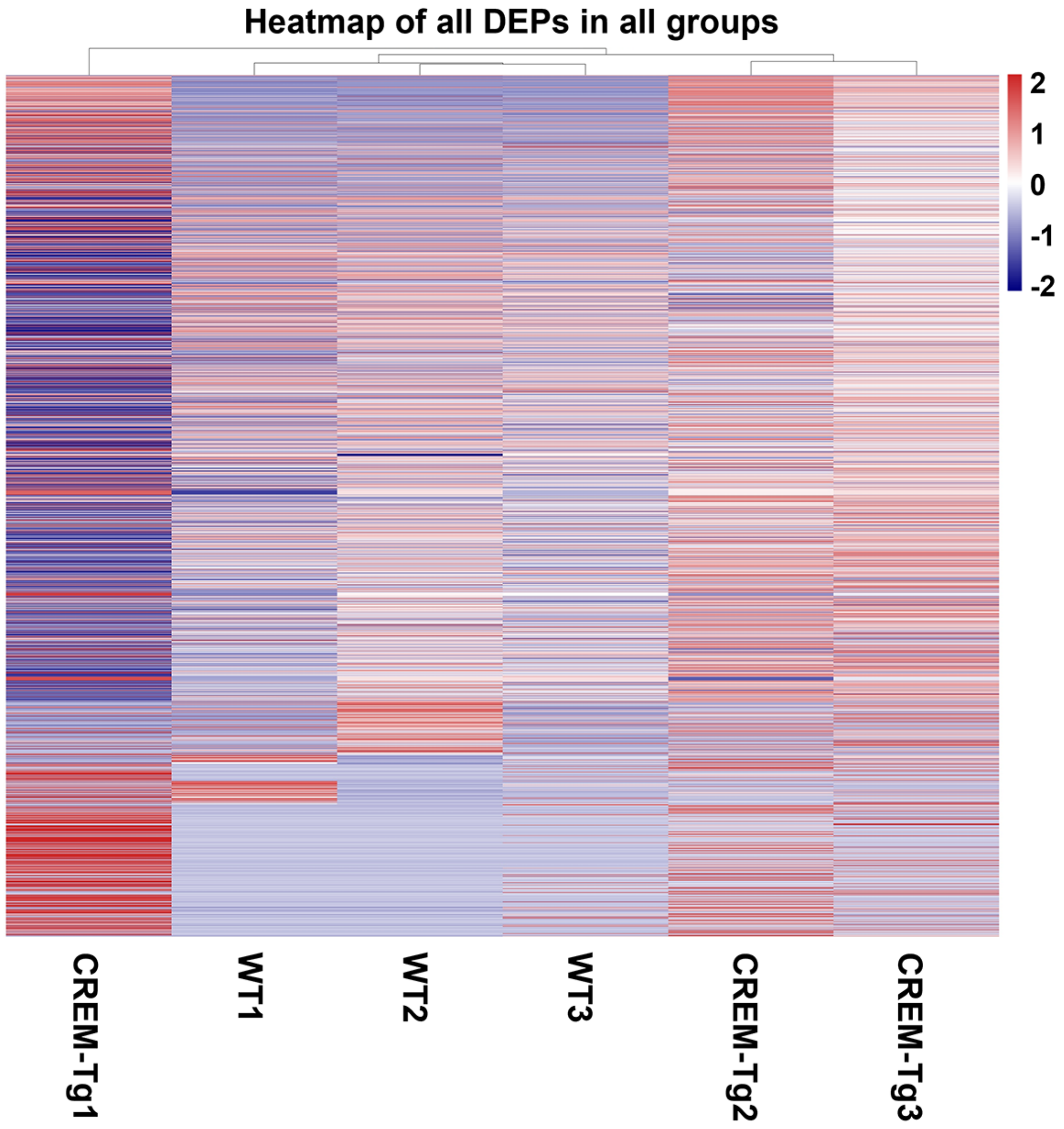

**Supplemental Figure 2. The tree plot of gene ontology (GO) term 'biological process' (BP; GO:0008150)**. Similar BPs are grouped by clusters. Dot size indicates the number of proteins in each process and dot color represents the protein expression fold change. Adjusted P values were calculated using the Benjamini-Hochberg procedure.

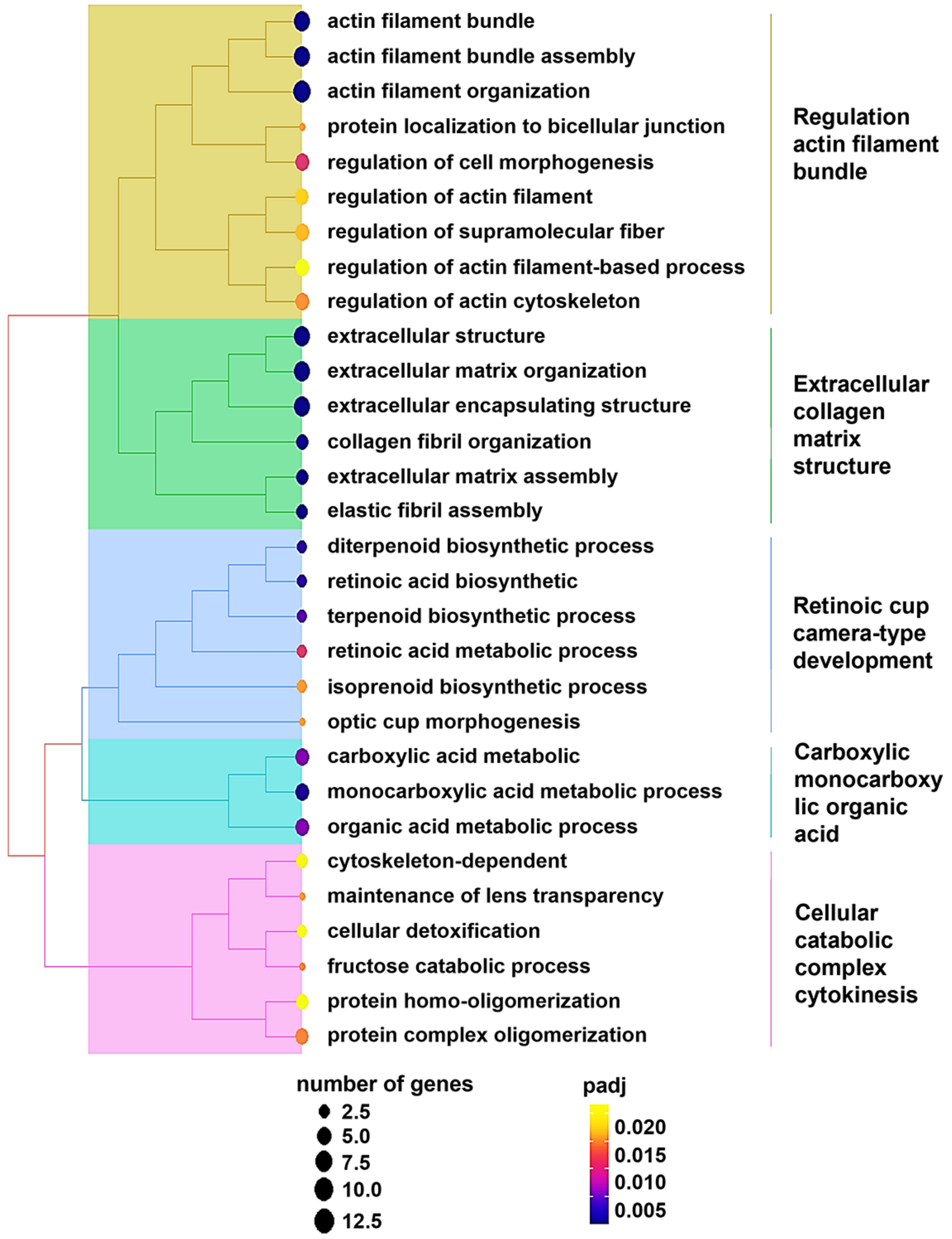

**Supplemental Figure 3. Cellular distribution of ITGAV.** The DISCO database was used to annotate the single-cell RNA sequencing data of cells from the heart tissue. The color darkness corresponds to the relative expression levels of integrin subunit alpha V (ITGAV) in specific cell subsets.

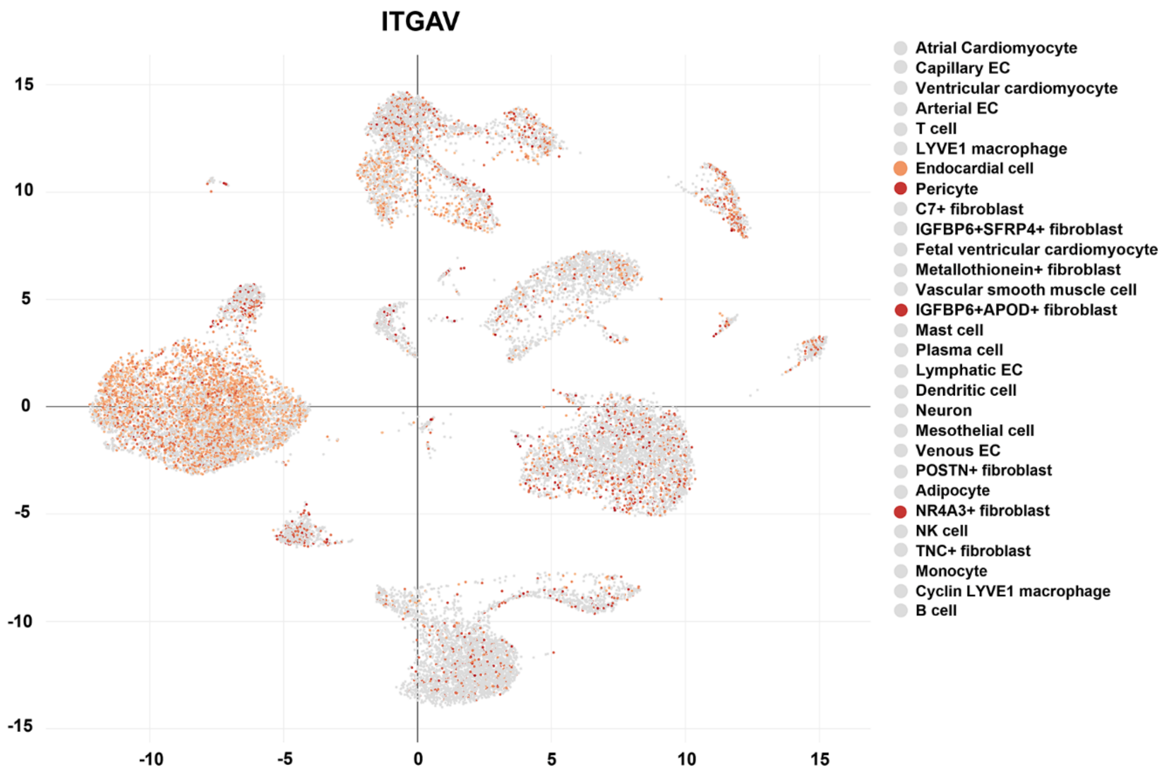

**Supplemental Figure 4. Cellular distribution of FBLN5.** The DISCO database was employed to annotate the single-cell RNA sequencing data of cells from the heart tissue. The color darkness corresponds to the high expression levels of fibulin 5 (FBLN5) in specific cell subsets.

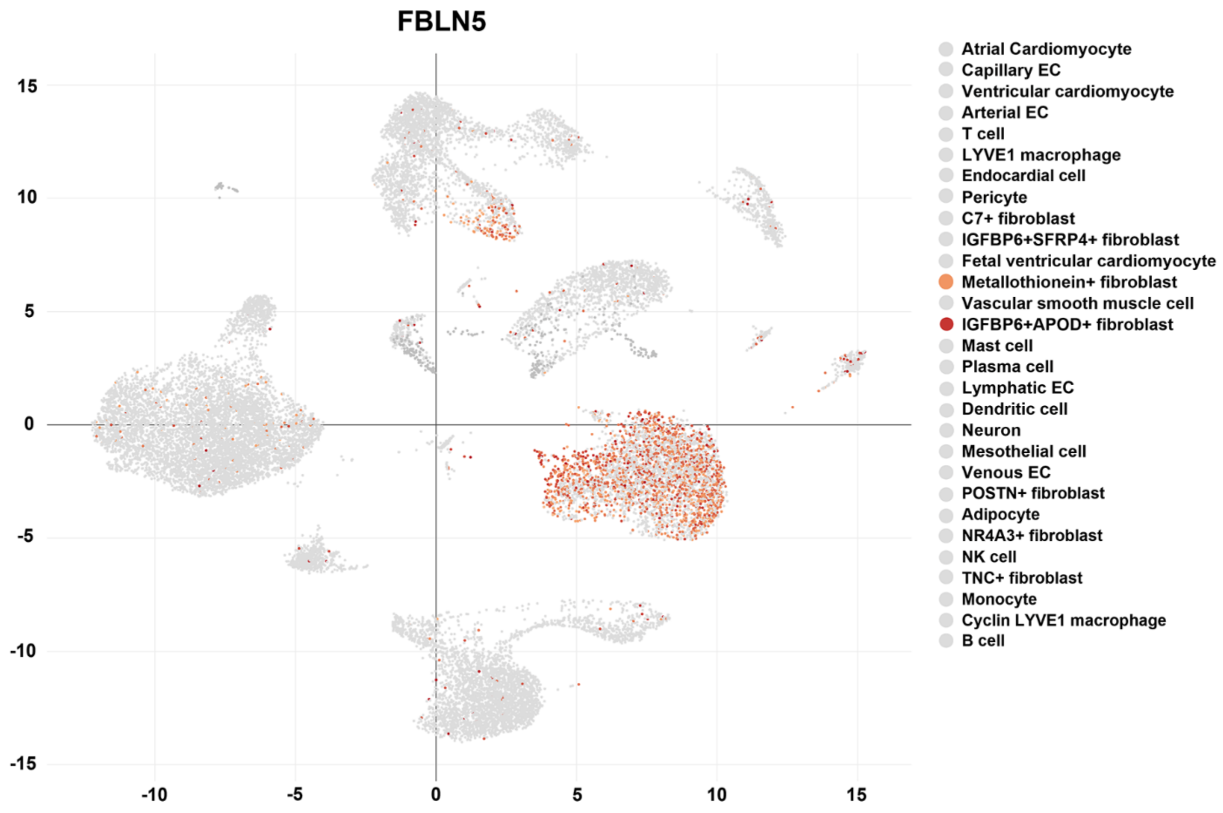

**Supplemental Figure 5. Cellular distribution of** LCP1**.** The DISCO database was employed to annotate the single-cell RNA sequencing data of cells from the heart tissue. The color darkness corresponds to the high expression levels of lymphocyte cytosolic protein 1 (LCP1) in specific cell subsets.

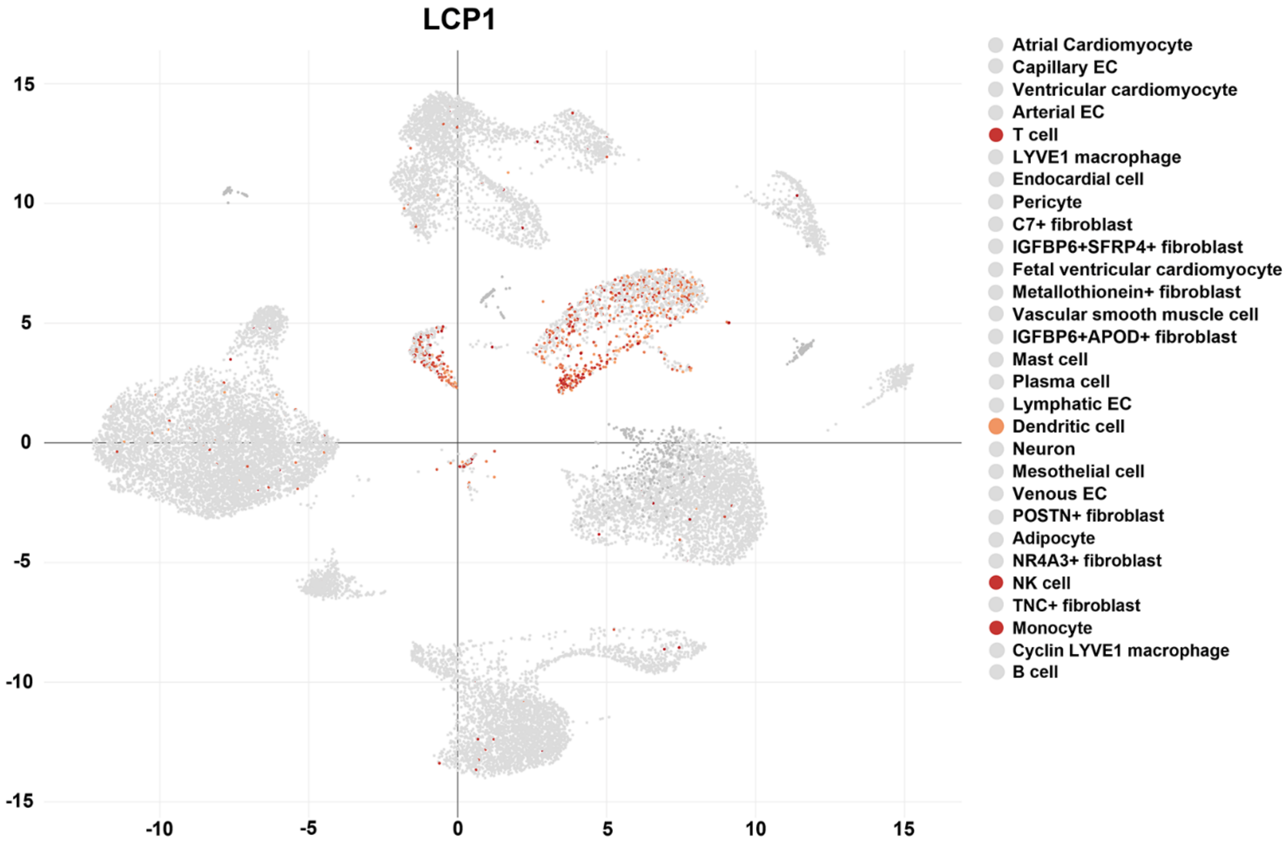
